# Temperature-sensitive cytoplasmic incompatibility across divergent *Wolbachia* partly reflects *cifB* transcription, not endosymbiont density

**DOI:** 10.64898/2026.03.31.715596

**Authors:** Basabi Bagchi, Lore Van Vlaenderen, Tim Wheeler, Elena Provencal, William R. Conner, Kyle McGuire, Brandon S. Cooper, J. Dylan Shropshire

## Abstract

Maternally transmitted *Wolbachia* bacteria are common in insects, with many strains altering host reproduction through cytoplasmic incompatibility (CI). CI kills embryos fertilized by *Wolbachia*-bearing males unless those embryos also carry *Wolbachia*, which favors females with *Wolbachia* and drives the endosymbiont to higher frequencies in host populations. Strong CI now underpins successful applications that rely on maintaining pathogen-blocking *Wolbachia* transinfections in vector populations to reduce arboviral disease transmission. Temperature modulates CI strength (the proportion of embryos killed), with consequences for *Wolbachia* prevalence in natural and transinfected populations. Yet the mechanisms regulating temperature-sensitive CI-strength variation are poorly understood. We quantified CI strength across eight divergent *Drosophila*-associated *Wolbachia* strains at four temperatures (18°C–26°C), while characterizing development time, *Wolbachia* and *Wovirus* densities, and transcription of the CI-inducing gene *cifB*. Four of eight *Wolbachia* strains exhibited temperature-sensitive CI, three of which induced CI at multiple temperatures. Of these three, two expressed significantly more *cifB* at the temperature yielding stronger CI, whereas testes *Wolbachia* density did not predict CI strength. Notably, *cifB*-transcript levels were consistently decoupled from *Wolbachia* and *Wovirus* densities, suggesting that *cifB* transcription is not regulated solely by symbiont abundance. We also report temperature-sensitive rescue of CI, *Wolbachia*-associated developmental acceleration, and strain-specific *Wovirus*-*Wolbachia* covariance. Our findings reveal temperature as a pervasive modulator of *Wolbachia*-host interactions at multiple levels and extend evidence that *cifB* transcription partly predicts variable CI strength across strain identities, male ages, and now temperatures. CI variation unaccounted for by *cifB* transcription points toward additional regulatory or post-transcriptional mechanisms that we discuss.

## Introduction

Heritable endosymbiotic bacteria are pervasive, influencing the physiology, ecology, and evolution of their arthropod hosts (Moran et al. 2008; Kaur et al. 2021; McCutcheon 2021; Hoffmann and Cooper 2024). Because temperature fundamentally shapes the physiology and fitness of ectotherms (Huey and Stevenson 1979; Clarke 1993; Angilletta 2009; Somero et al. 2017), it is a critical but underexplored variable for understanding endosymbiont-host interactions. Temperature can affect endosymbiont densities in host tissues (Dunbar et al. 2007; Anbutsu et al. 2008; Burke et al. 2010; Fan and Wernegreen 2013; Shan et al. 2014; Ross et al. 2017; Ross et al. 2020), alter endosymbiont-mediated phenotypes (Doremus et al. 2018; Ross et al. 2019a; Higashi et al. 2020; Corbin et al. 2021; Holliman et al. 2025), and modulate the fidelity of maternal endosymbiont transmission (Anbutsu et al. 2008; Osaka et al. 2008; Ross et al. 2017; Hague et al. 2020b). Some endosymbionts (*e.g., Wolbachia*) also modify host thermal preferences (Truitt et al. 2018; Hague et al. 2020a; Strunov et al. 2023), which can feed back on the expression of other temperature-sensitive traits (Dillon et al. 2009), including those that govern endosymbiont prevalence (Kriesner et al. 2016; Hague et al. 2022; Martins et al. 2023). Hence, understanding how temperature shapes endosymbiont-host dynamics is increasingly important as populations encounter novel thermal regimes (Corbin et al. 2017; Hector et al. 2022; Iltis et al. 2022).

*Wolbachia* bacteria are the most prevalent heritable endosymbionts, found in about half of terrestrial arthropod species (Weinert et al. 2015), though often at varied frequencies (Kriesner et al. 2013; Schuler et al. 2016; Cooper et al. 2017; Wheeler et al. 2021; Hague et al. 2022; Sanaei et al. 2022; Shastry et al. 2022). While a mix of vertical transmission (within host species) and horizontal transfer (between host species) contributes to *Wolbachia* prevalence across divergent hosts (O’Neill et al. 1992; Raychoudhury et al. 2009; Gerth and Bleidorn 2017; Turelli et al. 2018; Cooper et al. 2019; Vancaester and Blaxter 2023; Shropshire et al. 2026), cytoplasmic incompatibility (CI) is a key determinant of their spread within host populations (Hoffmann et al. 1990; Turelli and Hoffmann 1991; Kriesner et al. 2013). CI kills embryos fertilized by *Wolbachia*-bearing males unless rescued by maternal *Wolbachia* (Yen and Barr 1973; Shropshire et al. 2020). This conditional rescue confers a relative fitness advantage to *Wolbachia*-bearing females, whose offspring are compatible with males with and without *Wolbachia* (Hoffmann et al. 1990). The magnitude of this advantage depends on CI strength (*i.e*., the proportion of embryos killed in a CI cross), with strong CI driving *Wolbachia* to higher frequencies (Turelli and Hoffmann 1991; Hoffmann et al. 1996; Kriesner et al. 2013; Meany et al. 2019). CI also underpins *Wolbachia*-based applications that use strong CI to suppress vector and pest populations (Laven 1967; Mains et al. 2016; Zheng et al. 2019; Crawford et al. 2020) and drive pathogen-blocking *Wolbachia* transinfections into mosquito populations to reduce arboviral disease transmission (Hoffmann et al. 2011; Utarini et al. 2021; de Morais Batista et al. 2026). Despite CI’s importance, mechanisms governing CI-strength variation remain poorly understood.

What is clear is that *Wolbachia* genomes, host genotypes, host ontogeny, and the environment can shape CI strength (Poinsot et al. 1998; Reynolds and Hoffmann 2002; Veneti et al. 2003; Yamada et al. 2007; Bordenstein and Bordenstein 2011; Cooper et al. 2017; Layton et al. 2019; Hague et al. 2020b; Shropshire et al. 2021a; Shropshire et al. 2022; Ohata et al. 2025), with temperature being a particularly important environmental modulator. In *Ae. aegypti*, high rearing temperatures weaken CI, with thermal sensitivity varying among *Wolbachia* transinfections (Ross et al. 2017; Ross et al. 2019a; Gu et al. 2022; Duran-Ahumada et al. 2024). These effects have reduced transinfection frequencies in field populations following heatwaves (Ross et al. 2020), raising concerns about temperature effects on the efficacy of *Wolbachia*-based applications in some regions (Gu et al. 2022; Ross et al. 2023). Thermal sensitivity of CI induced by *Wolbachia* and other endosymbionts (Doremus et al. 2019; Doremus et al. 2020; Proctor et al. 2024) is taxonomically widespread, observed across flies (Wright and Wang 1980; Trpis et al. 1981; Hoffmann et al. 1986; Clancy and Hoffmann 1998), wasps (Bordenstein and Bordenstein 2011; Nasehi et al. 2022), and mites (van Opijnen and Breeuwer 1999; Lu et al. 2012). However, *Wolbachia*-induced CI seems temperature-resistant in some systems, including *Culex pipiens* mosquitoes (Sicard et al. 2021), *Leptopilina* wasps (Mouton et al. 2006), and *Tribolium* beetles (Gharabigloozare and Bleidorn 2022).

While the mechanisms underlying temperature-sensitive CI-strength variation remain unknown, there are several hypotheses. The *Wolbachia* density hypothesis is well established, positing that higher densities in testes produce stronger CI. Support for this hypothesis is mixed. Warm temperatures often reduce *Wolbachia* densities and weaken CI, as shown in *Drosophila simulans* flies (Clancy and Hoffmann 1998), *Habrobracon hebetor* wasps (Nasehi et al. 2022), and transinfected *Ae. aegypti* (Ross et al. 2017). In contrast, warm temperatures lower *Wolbachia* densities without affecting CI strength in *Leptopilina* and *Tribolium* (Mouton et al. 2006; Gharabigloozare and Bleidorn 2022); and in *Nasonia vitripennis* wasps, both warm and cold temperatures reduce *Wolbachia* densities, but CI is weaker under warm and stronger under cold temperatures (Bordenstein and Bordenstein 2011). A related hypothesis, the *cifB*-dosage hypothesis, shifts focus to the molecular level. CI is caused by *cifA* and *cifB* genes encoded by prophage WO (*Wovirus*) (Beckmann et al. 2017; LePage et al. 2017; Shropshire et al. 2018; Shropshire and Bordenstein 2019). *cifB*-transcript levels generally predict CI-strength variation across divergent *Wolbachia* (Shropshire et al. 2022) and sometimes predict age-sensitive variation in CI strength (Shropshire et al. 2021a). Whether temperature modulates *cifB* expression is largely unexplored, though in *H. hebetor*, high temperatures weakened CI despite increasing *cifB* transcription (Nasehi et al. 2022). Temperature could also modulate CI through indirect mechanisms. For instance, prolonging ectotherm development under cool conditions may extend the window for CI factors to act during spermatogenesis (Doremus et al. 2019; Doremus et al. 2020; Shropshire et al. 2022). Additionally, temperature stress may induce *Wovirus* lytic activity, which would reduce *Wolbachia* densities and potentially *cifB* transcription indirectly (Bordenstein and Bordenstein 2011; Nasehi et al. 2022). These non-mutually exclusive hypotheses suggest a multifactorial basis for temperature-sensitive CI and motivate comparisons of CI strength and its candidate predictors across systems and temperatures.

Here, we test whether variable CI strength across temperatures can be explained by molecular, organismal, and developmental factors using a comparative approach across eight *Drosophila*-associated *Wolbachia* strains diverged about 7.5 million years ago (MYA) (Shropshire et al. 2026). Of these, only *w*Ri of *D. simulans* has previously been characterized for temperature-sensitive CI (Hoffmann et al. 1986; Clancy and Hoffmann 1998); and while *w*Mel exhibits well-documented thermal sensitivity in transinfected *Ae. aegypti* (Ross et al. 2017; Ross et al. 2019a), it has not been characterized in its native *D. melanogaster* host, leaving seven of our eight systems unexplored. We reared males at four temperatures (18°C, 20°C, 23°C, and 26°C) and quantified CI strengths, development times, *Wolbachia* densities, *Wovirus* dynamics, and *cifB*-transcript levels. This framework allowed us to identify factors that predict CI strength across focal strains versus those with strain-specific effects, and to test whether *cifB* transcription predicts temperature-sensitive CI — a relationship not previously examined beyond *H. hebetor* (Nasehi et al. 2022). We show that temperature effects on CI are strain-specific at both organismal and molecular levels, with *cifB* transcription emerging as the best predictor of temperature-sensitive CI strength. Notably, *cifB*-transcript levels are decoupled from *Wolbachia* densities, while *Wovirus* and *Wolbachia* densities covary in strain-specific patterns across temperatures. We also report temperature-sensitive rescue of CI and *Wolbachia*-associated developmental acceleration. Together, our findings add to a growing body of literature (*e.g*., Bordenstein and Bordenstein 2011; Murdock et al. 2014; Ross et al. 2017; Ross et al. 2019a; Hague et al. 2020a; Chrostek et al. 2021; Hague et al. 2022; Nasehi et al. 2022; Caragata 2023) supporting temperature as a modulator of *Wolbachia*-host interactions. We discuss these findings and our *cifB*-transcription results, which support that additional regulatory or post-transcriptional mechanisms contribute to temperature-sensitive variation in CI strength.

## Results

### Focal *Wolbachia* diverged about 7.5 MYA and occupy hosts sampled from thermally diverse regions

The eight *Wolbachia* strains in our study diverged approximately 7.5 MYA [conservative plausible range (CPR): 1.4 to 22 MYA] (Shropshire et al. 2026) and comprise two clades: one that includes four “*w*Mel-like” variants (*w*Mel, *w*Tei, *w*Seg, *w*Cha) plus *w*Bic, and another that includes two “*w*Ri-like” variants (*w*Ri, *w*Tri) plus *w*Ha (**Fig 1A; Fig S1**). *w*Mel-like variants are estimated to have diverged about 206 thousand to 2.4 MYA (Shropshire et al. 2026), while very closely related *w*Ri-like variants diverged about 14 to 218 thousand years ago (Turelli et al. 2018; Shropshire et al. 2026). *Drosophila* hosts that carry our focal *Wolbachia* diverged about 23 MYA (Suvorov et al. 2022) and were sampled from seven locations with distinct thermal profiles (**Fig 1B**). They included temperate climates such as California (*w*Ri in *D. simulans*) and Japan (*w*Tri in *D. triauraria*), equatorial regions including Cameroon (*w*Seg in *D. seguyi*) and Bioko (*w*Tei in *D. teissieri*), and other tropical and subtropical origins (*w*Ha, *w*Cha, and *w*Bic). While the exact sampling site of the *w*Mel-*D. melanogaster* genotype is unknown, *D. melanogaster* is globally distributed (Sprengelmeyer et al. 2020), with temperature contributing to observed phenotypic and genomic clines (David et al. 1977; Hoffmann et al. 2002; Hoffmann and Weeks 2007; Keller 2007; Schmidt and Paaby 2008; Adrion et al. 2015). Detailed geographic temperature analyses and associated statistics are presented in the **Supporting Results** (**Fig S2**) to illustrate the diversity of thermal environments across the sampled locations. In summary, our study spans significant *Wolbachia* strain divergence, with *Wolbachia-Drosophila* genotypes sampled from thermally diverse locations.

**Figure 1.**
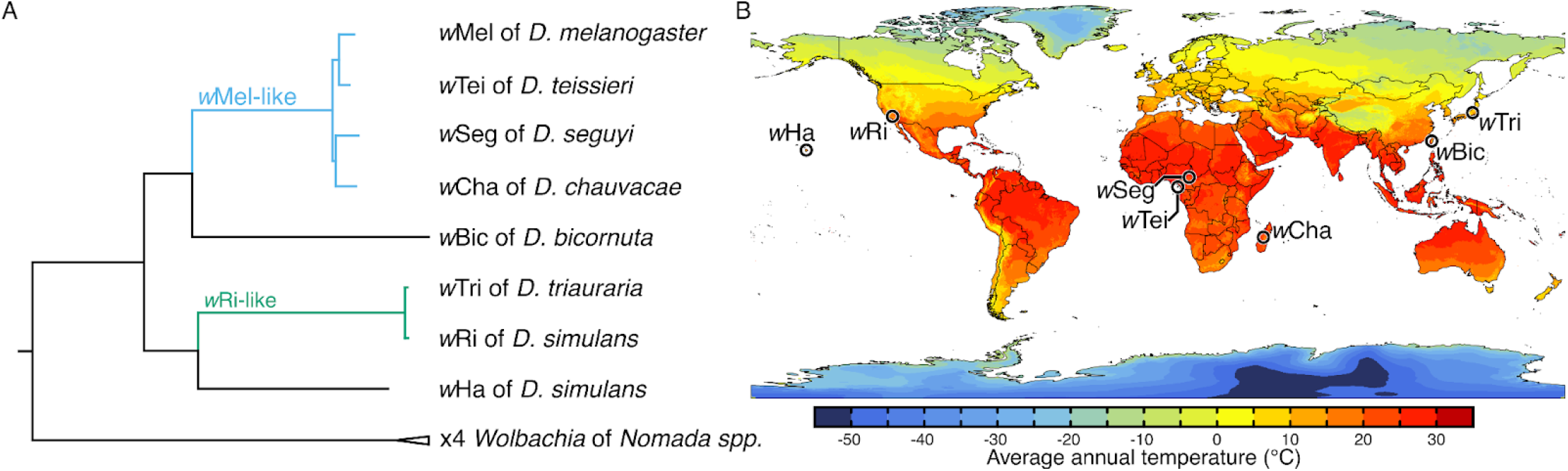
Phylogenetic relationships of the focal *Drosophila-associated Wolbachia* and sites of genotype sampling. **(A)** A phylogram based on 331 genes (285,540 bp). Four *Wolbachia* [(((*w*Nfa,*w*Nleu),wNpa),*w*Nfe)] from *Nomada* bee species were included as an outgroup to the eight focal *Wolbachia* strains associated with *Drosophila*. All *Wolbachia* belong to supergroup A and include closely related *w*Mel-like (*w*Mel, *w*Tei, *w*Seg, and *w*Cha) and *w*Ri-like (*w*Tri and *w*Ri) *Wolbachia*, as well as *w*Bic and *w*Ha. In **Fig S1** we present a cladogram placing novel *w*Cha among several known *w*Mel-like variants. The eight *Drosophila*-associated variants diverged approximately 1.4 to 22 MYA according to the estimates from Shropshire et al. (2026). Branch lengths are proportional to estimated substitutions per site. All nodes had posterior probabilities of 1. See **Table S1** for genome accession numbers. **(B)** Approximate sampling sites (black circles) for each *Wolbachia-Drosophila* genotype overlaid on a map displaying average annual temperatures (1970–2000) from WorldClim 2.1 data (Hijmans et al. 2005). The geographic origin for the *w*Mel-*D. melanogaster* genotype is unknown. See **Fig S2** for the average monthly temperatures by country or US state.

### Temperature modulated CI and CI rescue

To determine the effects of temperature on CI strength, we reared males from egg-to-adult at four experimental temperatures (18°C, 20°C, 23°C, and 26°C), crossed them to virgin females (maintained at 23°C) at each experimental temperature, and quantified egg hatchability. Due to rearing constraints, we collected males with and without *w*Tei at only three temperatures (20°C, 23°C, and 26°C) and males with and without *w*Tri at only two temperatures (23°C and 26°C). This reflects that our coolest temperature was not suitable for rearing either host species, and that 20°C was too cool to rear *D. triauraria*. We report results from three cross types: aposymbiotic males × aposymbiotic females (compatible control), symbiotic males × aposymbiotic females (CI), and symbiotic males × symbiotic females (rescue).

#### Thermal effects on embryo hatching depended on cross type and differ among systems

To model variation in embryo hatching across strains, temperatures, and cross types, we used a zero-inflated binomial generalized linear mixed model (GLMM; *N* = 1,339 crosses and 32,695 embryos; **Fig S3**). The model included symbiont-host system (eight focal systems), temperature (18°C, 20°C, 23°C, and 26°C), and cross type (compatible, CI, and rescue) as fixed effects with all two-way and three-way interactions, and an observation-level random effect. In order of decreasing effect size (η^2^_p_), significant terms included: cross type (η^2^_p_ = 0.46, χ^2^_2_ = 1125.85, *P* = 3.3e-245), system × cross type (η^2^_p_ = 0.10, χ^2^_14_ = 145.88, *P* = 4.8e-24), system × temperature × cross type (η^2^_p_ = 0.08, χ^2^_36_ = 120.13, *P* = 5.5e-11), system × temperature (η^2^_p_ = 0.07, χ^2^_18_ = 94.94, *P* = 1.9e-12), system (η^2^_p_ = 0.03, χ^2^_7_ = 42.29, *P* = 4.6e-7), and temperature (η^2^_p_ = 0.01, χ^2^_3_ = 16.08, *P* = 1.1e-3). The temperature × cross type interaction was not significant (η^2^_p_ = 0.003, χ^2^_6_ = 3.64, *P* = 0.72), indicating no overall temperature effect specific to cross type when averaged across systems. However, the significant three-way interaction (system × temperature × cross) demonstrates that temperature effects on cross-type-specific hatch rates differ among systems. Together, these results indicate that cross type is the dominant predictor of hatching success, while temperature effects on egg hatch are system-dependent.

#### Seven of eight Wolbachia induced CI in all thermal treatments

To test for CI at each temperature, we computed contrasts from the GLMM comparing CI crosses to compatible crosses within each strain × temperature combination. These contrasts yield odds ratios that quantify CI strength (*OR*_CI_; 1 = no CI, < 1 = CI). Seven of eight strains exhibited *OR*_CI_ significantly below 1 at all tested temperatures (*P* < 0.05; **Fig S4**), indicating consistent CI induction across thermal conditions. Across all strains and temperatures, *w*Cha at 23°C produced the strongest CI (*OR*_CI_ = 2.9e-5 [6.9e-7 to 1.3e-3], *P* = 5.3e-8). At 18°C (*OR*_CI_ = 6.1e-5 [7.5e-6 to 4.9e-4], *P* = 8.7e-20), 20°C (*OR*_CI_ = 5.4e-5 [1.2e-5 to 2.4e-4], *P* = 2.4e-38), and 26°C (*OR*_CI_ = 8.8e-5 [1.9e-5 to 4.0e-4], *P* = 1.5e-33), *w*Ri caused the strongest CI. Conversely, *w*Mel caused the weakest CI at 18°C (*OR*_CI_ = 0.024 [5.6e-3 to 0.1], *P* = 4.2e-7), 20°C (*OR*_CI_ = 7.1e-3 [1.9e-3 to 0.027], *P* = 3.5e-13), 23°C (*OR*_CI_ = 0.12 [0.034 to 0.44], *P* = 1.4e-3), and 26°C (*OR*_CI_ = 0.037 [9.4e-3 to 0.14], *P* = 1.9e-6). *w*Tei did not cause CI at 20°C (*OR*_CI_ = 0.87 [0.099 to 7.54], *P* = 0.90) and 26°C (*OR*_CI_ = 1.27 [0.11 to 14.25], *P* = 0.84), though low compatible-cross hatch rates at these temperatures (see Discussion) limit power to detect CI. *w*Tei caused moderate CI at 23°C (*OR*_CI_ = 4.8e-3 [3.9e-4 to 0.059], *P* = 3.0e-5; **Fig S4**). The temperature sensitivity of *w*Tei CI may have contributed to variable CI penetrance observed among studies (Zabalou et al. 2004; Zabalou et al. 2008; Martinez et al. 2015; Cooper et al. 2017), in addition to well-characterized host background effects (Cooper et al. 2017). In summary, all eight *Wolbachia*-host systems show CI in at least one temperature, though CI strength varies across systems and temperatures.

#### Four of eight Wolbachia exhibited temperature-sensitive CI-strength variation

Having established that all eight *Wolbachia* induce CI, we next tested whether temperature modulates CI strength. Since compatible cross hatch rates vary with temperature in several systems (see **Supporting Results**; **Fig S3**), we fitted strain-specific binomial GLMMs including only CI and compatible crosses, with a cross × temperature interaction term. The cross effect captured the difference between crosses, and the interaction term tested whether differences varied with temperature, while accounting for CI-independent variation in hatching. We calculated effect sizes from the interaction term, quantifying the magnitude of temperature effects on CI strength. Five strains showed significant overall temperature effects on CI strength: *w*Tei (η^2^ = 0.13, χ^2^_2_ = 7.27, *P* = 0.026), *w*Mel (η^2^ = 0.09, χ^2^_3_ = 16.9, *P* = 7.5e-4), *w*Ri (η^2^ = 0.08, χ^2^_3_ = 17.4, *P* = 5.8e-4), *w*Bic (η^2^ = 0.07, χ^2^_3_ = 8.56, *P* = 0.036), and *w*Ha (η^2^ = 0.06, χ^2^_3_ = 11.1, *P* = 0.011). Three strains showed no significant overall temperature effect: *w*Cha (η^2^ = 0.09, χ^2^_3_ = 7.46, *P* = 0.059), *w*Tri (η^2^ = 0.06, χ^2^_1_ = 1.91, *P* = 0.167), and *w*Seg (η^2^ = 0.03, χ^2^_3_ = 3.52, *P* = 0.318).

To identify differences in CI strength between specific temperatures, we performed pairwise temperature contrasts within each strain from the overall GLMM (which included all strains, temperatures, and their interactions), yielding *OR*_CI,T_ values (*OR*_CI,T_ = *OR*_CI,cool_ / *OR*_CI,warm_; > 1 stronger CI at warm, < 1 = stronger CI at cool). Four strains showed significant pairwise differences (**Fig 2**). *w*Mel produced stronger CI at 20°C than at 23°C (*OR*_CI,T_ = 0.059 [4.9e-3 to 0.70], *P* = 0.016), while other comparisons were not significant. *w*Tei showed CI only at 23°C, which was significantly stronger than at 20°C (*OR*_CI,T_ = 180 [3.16 to 1.0e+4], *P* = 3.1e-3) and 26°C (*OR*_CI,T_ = 3.8e-3 [5.4e-5 to 0.26], *P* = 3.1e-3), where we did not observe significant CI. *w*Ri produced the weakest CI at 23°C, relative to 18°C (*OR*_CI,T_ = 0.033 [1.2e-3 to 0.88], *P* = 0.016), 20°C (*OR*_CI,T_ = 0.029 [2.1e-3 to 0.40], *P* = 2.3e-3), and 26°C (*OR*_CI,T_ = 21.1 [1.47 to 301.5], *P* = 7.5e-3) where CI strength was similar. For *w*Ha, CI was weaker at 20°C than at 23°C (*OR*_CI,T_ = 25.9 [1.18 to 566.3], *P* = 0.016) and 26°C (*OR*_CI,T_ = 32.39 [1.62 to 649.6], *P* = 0.013), with intermediate CI strength observed at 18°C. In contrast, *w*Seg, *w*Cha, *w*Bic, and *w*Tri showed no significant pairwise differences in CI strength between temperatures (all *P* > 0.05; **Fig 2**).

**Figure 2.**
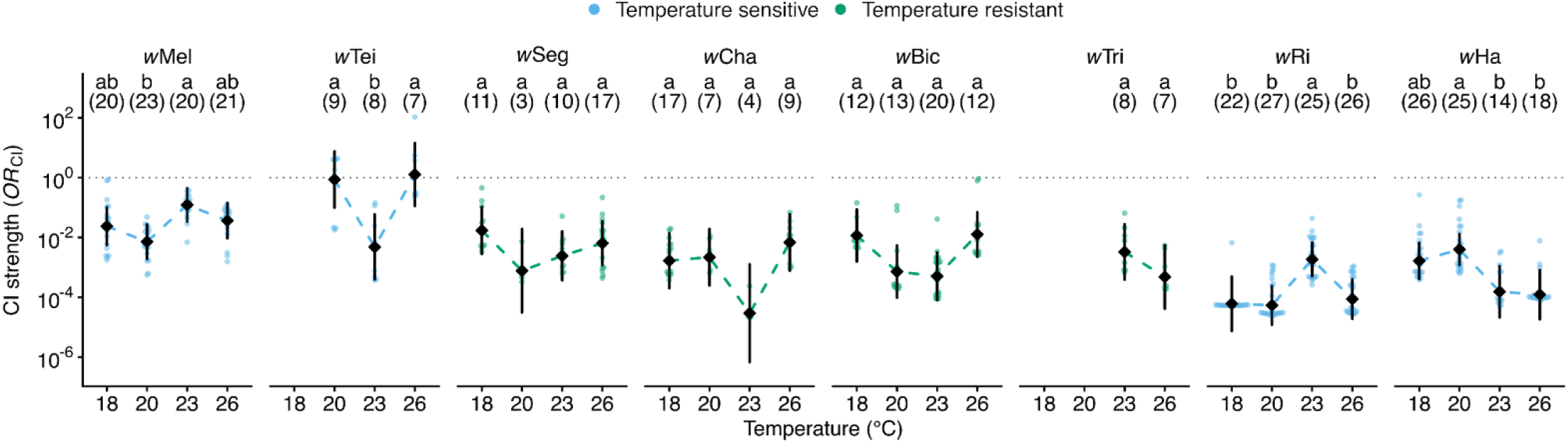
CI strength is modulated by temperature in a strain-specific manner. Temperature modulated CI strength in four of eight focal *Wolbachia*-host symbioses (*w*Mel, *w*Tei, *w*Ri, *w*Ha). CI strength was quantified as the odds ratio of embryo hatching in CI crosses relative to compatible crosses (*OR*_CI_; 1 = no CI, < 1 = CI). Dotted horizontal lines highlight *OR*_CI_ = 1. Diamonds represent model-estimated odds ratios per strain and temperature, with 95% confidence intervals as vertical lines. Individual points show model-predicted odds ratios for each CI cross replicate, derived from binomial GLMM-predicted hatch odds relative to the compatible cross baseline. Dashed lines connect estimated means across temperatures to highlight thermal response patterns. Letters indicate significant differences in CI strength observed among temperatures within each strain (FDR-corrected, α = 0.05); shared letters denote non-significant differences. Sample sizes per group are in parentheses.

Based on the combined evidence, we classified *w*Mel, *w*Tei, *w*Ri, and *w*Ha CI as temperature-sensitive and *w*Seg, *w*Cha, *w*Bic, and *w*Tri as temperature-resistant. *w*Bic showed a significant overall temperature effect but no significant pairwise differences, suggesting modest but distributed effects. We conservatively classified *w*Bic as temperature-resistant. Notably, the temperature-sensitive and temperature-resistant classes each included *w*Mel-like and *w*Ri-like *Wolbachia* variants. Our findings indicate that temperature effects on CI strength are system-dependent, including among closely related *Wolbachia* strains.

### Three Wolbachia strains exhibited temperature-sensitive CI-rescue efficiency

While our primary goal was to understand temperature-sensitive CI-strength variation, we also tested whether CI rescue efficiency showed similar temperature sensitivity since it reflects the ability of *Wolbachia*-bearing females to restore compatibility in CI crosses. We tested rescue efficiency by contrasting rescue and compatible crosses within the GLMM (*OR*_R_; 1 = complete rescue, < 1 = incomplete). Because all females were maintained at 23°C until crossing, temperature effects on rescue reflect crossing conditions and male developmental effects. *w*Mel, *w*Tei, *w*Seg, and *w*Tri all exhibited complete rescue across all tested temperatures (*P* > 0.05; **Fig S4**). In contrast, *w*Bic at 23°C (*OR*_R_ = 0.15 [0.024 to 0.96], *P* = 0.045) and *w*Ha at 23°C (*OR*_R_ = 0.17 [0.029 to 0.97], *P* = 0.047; **Fig S4**), showed marginally incomplete rescue. *w*Cha showed incomplete rescue at 20°C (*OR*_R_ = 0.021 [2.5e-3 to 0.18], *P* = 4.8e-4; **Fig S4**), while *w*Ri was the only strain with incomplete rescue at multiple temperatures: 20°C (*OR*_R_ = 0.10 [0.014 to 0.43], *P* = 2.0e-3) and 26°C (*OR*_R_ = 0.07 [0.014 to 0.36], *P* = 1.4e-3; **Fig S4**). These results indicate that rescue efficiency, like CI strength, varies in a system-dependent manner with temperature, with closely related *w*Mel-like (*w*Mel, *w*Tei, *w*Seg versus *w*Cha) and *w*Ri-like (*w*Ri versus *w*Tri) differing in the temperature-sensitivity of CI rescue.

### Temperature influenced development time, but development time did not predict CI strength

#### Development time varied with temperature, system, and cytotype

In *Cardinium*-bearing *Encarsia* wasps, longer development time correlates with stronger CI (Doremus et al. 2019; Doremus et al. 2020). To test whether this pattern holds for *Wolbachia* across temperatures, we first analyzed development time (egg laying to first adult emergence) using PERMANOVA (*N* = 376). The model included system (six *Drosophila*-*Wolbachia* symbioses; *w*Ha and *w*Tri excluded due to insufficient sample sizes), cytotype (aposymbiotic and symbiotic), temperature (18°C, 20°C, 23°C, and 26°C), and all two- and three-way interactions. In order of decreasing variance explained, significant terms included: temperature (*R*^2^ = 0.64, *F*_3_ = 2458.5, *P* < 1.0e-4), system (*R*^2^ = 0.28, *F*_5_ = 643.2, *P* < 1.0e-4), system × temperature (*R*^2^ = 0.03, *F*_11_ = 33.8, *P* < 1.0e-4), system × cytotype × temperature (*R*^2^ = 0.01, *F*_11_ = 13.6, *P* < 1.0e-4), cytotype × temperature (*R*^2^ = 0.003, *F*_3_ = 10.4, *P* < 1.0e-4), system × cytotype (*R*^2^ = 0.003, *F*_5_ = 6.2, *P* < 1.0e-4), and cytotype (*R*^2^ = 0.002, *F*_1_ = 22.1, *P* < 1.0e-4). Hence, temperature and system identity explain significant variation in development time, with a smaller, yet statistically significant, effect of *Wolbachia* cytotype that varies across systems and temperatures.

#### Wolbachia accelerated development in wCha and wBic at specific temperatures

Temperature was the dominant factor influencing development time (see **Supporting Results**; **Fig S5**), but smaller, significant cytotype effects are also present. To further assess *Wolbachia* effects, we performed pairwise permutation tests comparing development times of aposymbiotic and symbiotic females within each strain-temperature combination (Δt; < 0 = symbiotic faster, > 0 = aposymbiotic faster). Four of six systems in this analysis exhibited no significant cytotype effect on development time at any temperature: *D. melanogaster* (largest effect at 18°C: Δt = −0.3 days, *P* = 0.84), *D. teissieri* (largest effect at 20°C: Δt = 0.6 days, *P* = 0.11), *D. seguyi* (23°C only: Δt = −0.44 days, *P* = 0.50), and *D. simulans* that carries *w*Ri (largest effect at 18°C: Δt = −0.28 days, *P* = 0.54; **Fig S6**). In contrast, *D. chauvacae* exhibited significantly faster development in *w*Cha-symbiotic vials at two temperatures: 20°C (Δt = −1.9 days, *P* = 0.002) and 26°C (Δt = −1.3 days, *P* = 0.05; **Fig S6**). *D. bicornuta* showed accelerated development in *w*Bic-symbiotic vials only at 18°C (Δt = −3.6 days, *P* = 0.011; **Fig S6**). Development times at 18°C for *D. chauvacae* and at 20°C, 23°C, and 26°C for *D. bicornuta* showed no significant differences between cytotypes (*P* > 0.05; **Fig S6**). These observations indicate that *Wolbachia* can accelerate development time in a strain-specific and temperature-sensitive manner.

#### Development time did not predict temperature-sensitive CI strength

Having characterized how development times varied with temperature and *Wolbachia* presence, we next tested whether development time predicted CI strength across temperatures, as observed in *Cardinium*-bearing *Encarsia* wasps (Doremus et al. 2019; Doremus et al. 2020). We used Bayesian phylogenetic mixed models to test whether development time predicts CI strength, yielding regression coefficients (β) and the probability of direction (*p*_d_), which indicates the certainty in the direction of β. We analyzed all strains with development time data (*w*Seg, *w*Tri, and *w*Ha excluded due to insufficient data), as well as subsets of strains with temperature-sensitive CI (*w*Mel, *w*Tei, *w*Ri) and temperature-resistant CI (*w*Cha, *w*Bic). Bayesian analysis of the relationship between CI strength and development time yielded small effects across all strains (β = −0.05 [−0.20 to 0.10], *R*^2^ = 0.45, *p*_d_ = 0.75, *n* = 19, 5 strains), strains with temperature-sensitive CI (β = −0.09 [−0.31 to 0.13], *R*^2^ = 0.64, *p*_d_ = 0.81, *n* = 11, 3 strains), and strains with temperature-resistant CI (β = 0.04 [−0.18 to 0.27], *R*^2^ = 0.21, *p*_d_ = 0.68, *n* = 8, 2 strains; **Fig S7**). Effect estimates were not credibly different from zero (all 95% credible intervals for β span zero) and directional certainty was weak (*p*_d_ = 0.68 to 0.81), providing no credible support for a relationship between development time and CI strength.

To complement cross-strain analyses, we examined within-strain concordance using Kendall’s τ, propagating uncertainty from estimated marginal means via Monte Carlo simulation. We report the rank correlation coefficient (τ) and the probability of direction (*p*_d_), the proportion of simulations agreeing with the sign of τ. Among the five strains analyzed, concordance patterns varied in direction and certainty: *w*Mel (τ = −0.31, *p*_d_ = 0.66, *n* = 4), *w*Ri (τ = −0.28, *p*_d_ = 0.61, *n* = 4), *w*Cha (τ = −0.07, *p*_d_ = 0.61, n = 4), *w*Bic (τ = +0.11, *p*_d_ = 0.82, *n* = 4), and *w*Tei (τ = +0.69, *p*_d_ = 0.70, *n* = 3). No strain showed complete concordance (τ = ±1), and *p*_d_ values indicated substantial uncertainty in the direction of within-strain relationships. These results reveal that development time does not reliably predict CI-strength variation across temperatures.

### Temperature modulated *Wolbachia* densities, but densities do not predict CI strength

#### Wolbachia densities varied by strain and thermal exposure

The *Wolbachia* density hypothesis posits that higher symbiont densities yield stronger CI (Breeuwer and Werren 1993). To test this hypothesis, we dissected testes from male siblings of flies used in CI experiments across temperatures, extracted genomic DNA, and quantified *Wolbachia* density relative to a conserved single-copy host gene (nAcRalpha-34E) via qPCR. We first describe how density varies with strain and temperature. We analyzed how *Wolbachia* densities varied with strain and temperature using a linear model on log_2_-transformed density values (*N* = 84 samples, each containing 5 pairs of testes). The model included *Wolbachia* strain (eight focal strains), temperature (18°C, 20°C, 23°C, and 26°C), and their interaction as fixed effects. According to effect sizes (ω^2^_p_), *Wolbachia* strain was the dominant predictor of density (ω^2^_p_ = 0.86, *F*_7_ = 75.5, *P* = 6.8e-26) with strain × temperature (ω^2^_p_ = 0.43, *F*_18_ = 4.7, *P* = 5.1e-6) and temperature (ω^2^_p_ = 0.28, *F*_3_ = 12.4, *P* = 2.6e-6) also contributing significantly. Therefore, strain identity primarily determines *Wolbachia* densities, while temperature effects on densities vary among strains.

#### Five Wolbachia exhibited temperature-sensitive Wolbachia densities

We next determined which strains exhibited temperature-sensitive densities by comparing estimated marginal means from the linear model across all pairwise temperature combinations within each strain, yielding risk ratios (*RR*; > 1 = higher density at cool, < 1 = higher density at warm). Effect sizes (ω^2^) quantify the magnitude of temperature’s effect per strain from strain-specific linear models. Three strains exhibited temperature-resistant *Wolbachia* densities, with no significant differences across any temperature comparisons: *w*Tei (ω^2^ = 0, largest effect: *RR*_20:23_ = 0.71 [0.25 to 2.02], *P* = 0.79), *w*Cha (ω^2^ = 0, largest effect: *RR*_18:26_ = 0.68 [0.21 to 2.17], *P* = 0.94), and *w*Tri (*RR*_23:26_ = 0.57 [0.24 to 1.33], *P* = 0.19; **Fig 3A**). In contrast, five strains displayed significant temperature sensitivity: *w*Ri (ω^2^ = 0.81), *w*Seg (ω^2^ = 0.69), *w*Mel (ω^2^ = 0.64), *w*Bic (ω^2^ = 0.59), and *w*Ha (ω^2^ = 0.39). *w*Ri density was lowest at cool temperatures, with 18°C similar to 20°C (*RR* = 0.51 [0.16 to 1.64], *P* = 0.15) and lower than 23°C (*RR* = 0.15 [0.047 to 0.49], *P* = 2.7e-4) and 26°C (*RR* = 0.20 [0.064 to 0.65], *P* = 0.001; **Fig 3A**). *w*Seg showed a similar thermal response, with density at 18°C increasing slightly at 20°C (*RR* = 0.36 [0.10 to 1.34], *P* = 0.057) and significantly at both 23°C (*RR* = 0.12 [0.032 to 0.43], *P* = 2.2e-4) and 26°C (*RR* = 0.20 [0.062 to 0.63], *P* = 0.001; **Fig 3A**). *w*Mel density was also lowest at 18°C and increased slightly at 20°C (*RR* = 0.58 [0.18 to 1.87], *P* = 0.25) and significantly at 23°C (*RR* = 0.24 [0.076 to 0.78], *P* = 0.005) and 26°C (*RR* = 0.18 [0.057 to 0.59], *P* = 0.001; **Fig 3A**). *w*Bic showed a distinct thermal response compared to other temperature-sensitive strains, with lower density at 26°C than at 18°C (*RR* = 3.02 [0.94 to 9.67], *P* = 0.018) and 20°C (*RR* = 9.87 [3.08 to 31.6], *P* = 9.4e-6); and higher density at 20°C than at 23°C (*RR* = 8.75 [2.73 to 28.0], *P* = 1.3e-5) and 18°C (*RR* = 0.31 [0.10 to 0.98], *P* = 0.015; **Fig 3A**). Finally, *w*Ha exhibited relatively modest temperature sensitivity, with lower density at 18°C than at 26°C (*RR* = 0.27 [0.084 to 0.86], *P* = 0.019); *Wolbachia* densities at intermediate temperatures were statistically similar to all other temperatures (*P* > 0.05; **Fig 3A**). Overall, focal *Wolbachia* exhibit diverse density responses to temperature: cold inhibition (*w*Mel, *w*Seg, *w*Ri, *w*Ha), an intermediate peak (*w*Bic), and temperature-resistance (*w*Tei, *w*Cha, *w*Tri). Again, closely related *Wolbachia* often differ in their responses to temperature.

**Figure 3.**
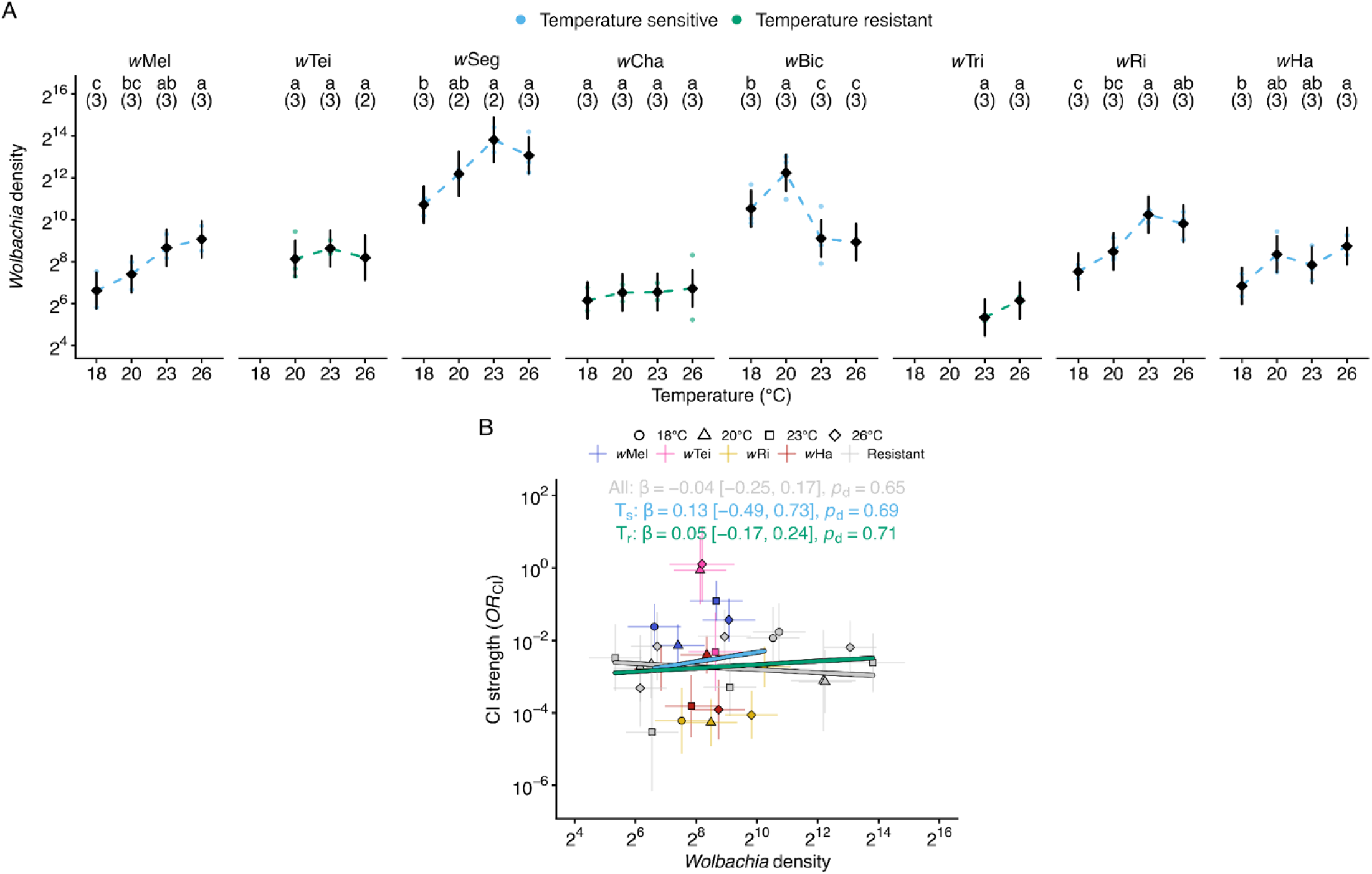
*Wolbachia* densities varied with temperature but did not predict CI strength. **(A)** *Wolbachia* densities in testes varied with temperature in five of eight focal systems (*w*Mel, *w*Seg, *w*Bic, *w*Ri, *w*Ha). Diamonds represent model-estimated mean *Wolbachia* densities per strain and temperature, with 95% confidence intervals as vertical lines. Individual points show *Wolbachia*-density values from biological replicates containing 5 pairs of testes. Dashed lines connect estimated means across temperatures to highlight thermal response patterns. Letters indicate significant differences in CI strength among temperatures within each strain (FDR-corrected, α = 0.05); shared letters denote non-significant differences. Sample sizes per group are in parentheses. **(B)** *Wolbachia* densities in testes did not significantly predict CI strength across strain-temperature combinations. Individual points represent estimated marginal means for *OR*_CI_ and *Wolbachia* density per strain and temperature combination, with 95% confidence intervals as vertical and horizontal lines. Regression lines and statistics (β [95% credible interval], *p*_d_) are derived from Bayesian phylogenetic mixed models fit separately to all strains (gray; *n* = 29, 8 strains), temperature-sensitive CI strains (T_s_; light blue; *n* = 15, 4 strains), and temperature-resistant CI strains (T_r_; green; *n* = 14, 4 strains). **(A, B)** *Wolbachia* densities were measured by qPCR as fold-change of *ftsZ* relative to host nAcRalpha-34E (primers: **Table S3**).

#### Wolbachia densities did not predict temperature-sensitive CI strength

With *Wolbachia* density variation characterized across strains and temperatures, we tested whether densities predict CI strength using Bayesian phylogenetic mixed models with random intercepts to account for evolutionary non-independence among strains. Bayesian analysis revealed no credible relationship between *OR*_CI_ and *Wolbachia* densities when considering all strains (β = −0.04 [−0.25 to 0.17], *R*^2^ = 0.43, *p*_d_ = 0.65, *n* = 29, 8 strains), strains with temperature-sensitive CI (*w*Mel, *w*Tei, *w*Ri, *w*Ha; β = 0.13 [−0.49 to 0.73], *R*^2^ = 0.62, *p*_d_ = 0.69, *n* = 15, 4 strains), or strains with temperature-resistant CI (*w*Seg, *w*Cha, *w*Bic, *w*Tri; β = 0.05 [−0.17 to 0.24], *R*^2^ = 0.15, *p*_d_ = 0.71, *n* = 14, 4 strains; **Fig 3B**). All credible intervals included zero and *p*_d_ values indicated little confidence in the direction of effects. Within-strain concordance analysis examined whether higher densities corresponded to stronger CI (more negative *OR*_CI_) across temperature pairs. In contrast to the *Wolbachia* density hypothesis, two strains showed confident positive concordance, where higher densities corresponded to weaker CI: *w*Ri (τ = +0.79, *p*_d_ = 0.97, *n* = 4) and *w*Mel (τ = +0.48, *p*_d_ = 0.95, *n* = 4). The remaining strains displayed weaker or uncertain concordance patterns: *w*Tei (τ = −0.79, *p*_d_ = 0.77, *n* = 3), *w*Tri (τ = −0.98, *p*_d_ = 0.80, *n* = 2), *w*Seg (τ = −0.43, *p*_d_ = 0.64, *n* = 4), *w*Cha (τ = +0.10, *p*_d_ = 0.68, *n* = 4), *w*Bic (τ = −0.21, *p*_d_ = 0.51, *n* = 4), and *w*Ha (τ = −0.26, *p*_d_ = 0.53, *n* = 4). Our findings indicate that, as with development time, *Wolbachia* densities are not a reliable predictor of CI-strength variation across temperatures.

### *Wovirus* dynamics covaried with temperature-sensitive *Wolbachia* densities

#### Wovirus density per Wolbachia varied by strain and thermal exposure

In *N. vitripennis*, temperature-sensitive *Wolbachia* (*w*VitA) densities correlate with *Wovirus* abundance, suggesting lytic prophage activity may drive *Wolbachia* density variation at stressful temperatures (Bordenstein and Bordenstein 2011). We quantified *Wovirus* densities — both per *Wolbachia* (to assess phage-bacterium dynamics) and per host (to capture total *Wovirus* load, which determines *cif*-gene copy number in host cells) — before examining how these metrics related to *Wolbachia* densities and CI strength. Using the same genomic DNA samples collected for *Wolbachia* density analyses, we first quantified *Wovirus* densities per *Wolbachia* (relative to the *Wolbachia* gene *ftsZ*) via qPCR. For strains containing multiple divergent prophage variants, density values were summed across all detected variants to approximate total *Wovirus* abundance. Across strains, only a single sr3WO variant from *w*Tei was not captured by our design. We analyzed how *Wovirus* densities vary with strain and temperature using a linear model on log_2_-transformed density values (*N* = 74 samples, each containing 5 pairs of testes). The model included *Wolbachia* strain (seven focal strains; *w*Bic excluded), temperature (18°C, 20°C, 23°C, and 26°C), and their interaction as fixed effects. *Wolbachia* strain was the dominant predictor of *Wovirus* densities per *Wolbachia* (ω^2^_p_ = 0.87, *F*_6_ = 86.0, *P* = 2.7e-24), with strain × temperature (ω^2^_p_ = 0.62, *F*_15_ = 9.0, *P* = 1.9e-9) and temperature (ω^2^_p_ = 0.40, *F*_3_ = 17.6, *P* = 6.9e-8) also contributing significantly. Our findings indicate that strain identity primarily determines *Wovirus* densities per *Wolbachia*, which also vary with temperature in a strain-specific manner.

#### Five Wolbachia exhibited temperature-sensitive Wovirus densities per Wolbachia

We next determined which strains exhibited temperature-sensitive *Wovirus* densities per *Wolbachia*. Two strains exhibited temperature-resistant *Wovirus* densities, with no significant differences across any temperature comparisons: *w*Tei (ω^2^ = 0.21, largest effect: *RR*_23:26_ = 1.39 [0.89 to 2.17], *P* = 0.17) and *w*Tri (*RR*_23:26_ = 1.08 [0.78 to 1.50], *P* = 0.62; **Fig 4A**). Five strains displayed significant temperature-sensitive *Wovirus* density variation. *w*Mel (ω^2^ = 0.96) *Wovirus* density at 18°C was similar to density at 20°C (*RR* = 0.86 [0.56 to 1.34], *P* = 0.37), but significantly lower than density at 23°C (*RR* = 0.26 [0.17 to 0.40], *P* = 2.2e-10) and 26°C (*RR* = 0.42 [0.27 to 0.66], *P* = 4.7e-6). Notably, *Wovirus* density at 26°C was lower than at 23°C (*RR* = 1.65 [1.06 to 2.57], *P* = 0.004; **Fig 4A**), indicating that the relationship between temperature and *Wovirus* density is non-monotonic in *w*Mel. *w*Ri (ω^2^ = 0.84) exhibited a similar thermal response pattern, with density at 18°C similar to 20°C (*RR* = 1.28 [0.83 to 2.00], *P* = 0.15) and lower than both 23°C (*RR* = 0.55 [0.36 to 0.86], *P* = 0.001) and 26°C (*RR* = 0.60 [0.39 to 0.93], *P* = 0.004; **Fig 4A**). *w*Seg and *w*Ha showed inverse thermal responses compared to *w*Mel and *w*Ri, with higher *Wovirus* density at cooler temperatures. In *w*Seg (ω^2^ = 0.29), *Wovirus* density at 18°C was similar to density at 20°C (*RR* = 1.25 [0.80 to 1.94], *P* = 0.21) and significantly higher than at 23°C (*RR* = 1.63 [1.04 to 2.53], *P* = 0.015) and 26°C (*RR* = 1.60 [1.03 to 2.49], *P* = 0.015; **Fig 4A**). For *w*Ha (ω^2^ = 0.42), density at 18°C was higher than at 20°C (*RR* = 2.13 [1.37 to 3.31], *P* = 1.3e-4), 23°C (*RR* = 1.56 [1.0 to 2.43], *P* = 0.012), and 26°C (*RR* = 1.6 [1.03 to 2.49], *P* = 0.015); whereas *Wovirus* densities from 20°C to 26°C were similar (*P* > 0.05; **Fig 4A**). *w*Cha (ω^2^ = 0.62) exhibited a distinct pattern where *Wovirus* density at 20°C was lower than at 18°C (*RR* = 1.8 [1.13 to 2.75], *P* = 0.005), 23°C (*RR* = 0.68 [0.44 to 1.06], *P* = 0.04), and 26°C (*RR* = 0.59 [0.38 to 0.92], *P* = 0.006), yet all other comparisons were similar (*P* > 0.05; **Fig 4A**). Overall, *w*Mel and *w*Ri tend to exhibit reduced *Wovirus* densities per *Wolbachia* at higher temperatures, *w*Seg and *w*Ha show increased *Wovirus* densities per *Wolbachia* at warmer temperatures, and *w*Cha displays a *Wovirus* density decline at intermediate temperatures. Again, patterns associated with closely related *Wolbachia* often differ.

**Figure 4.**
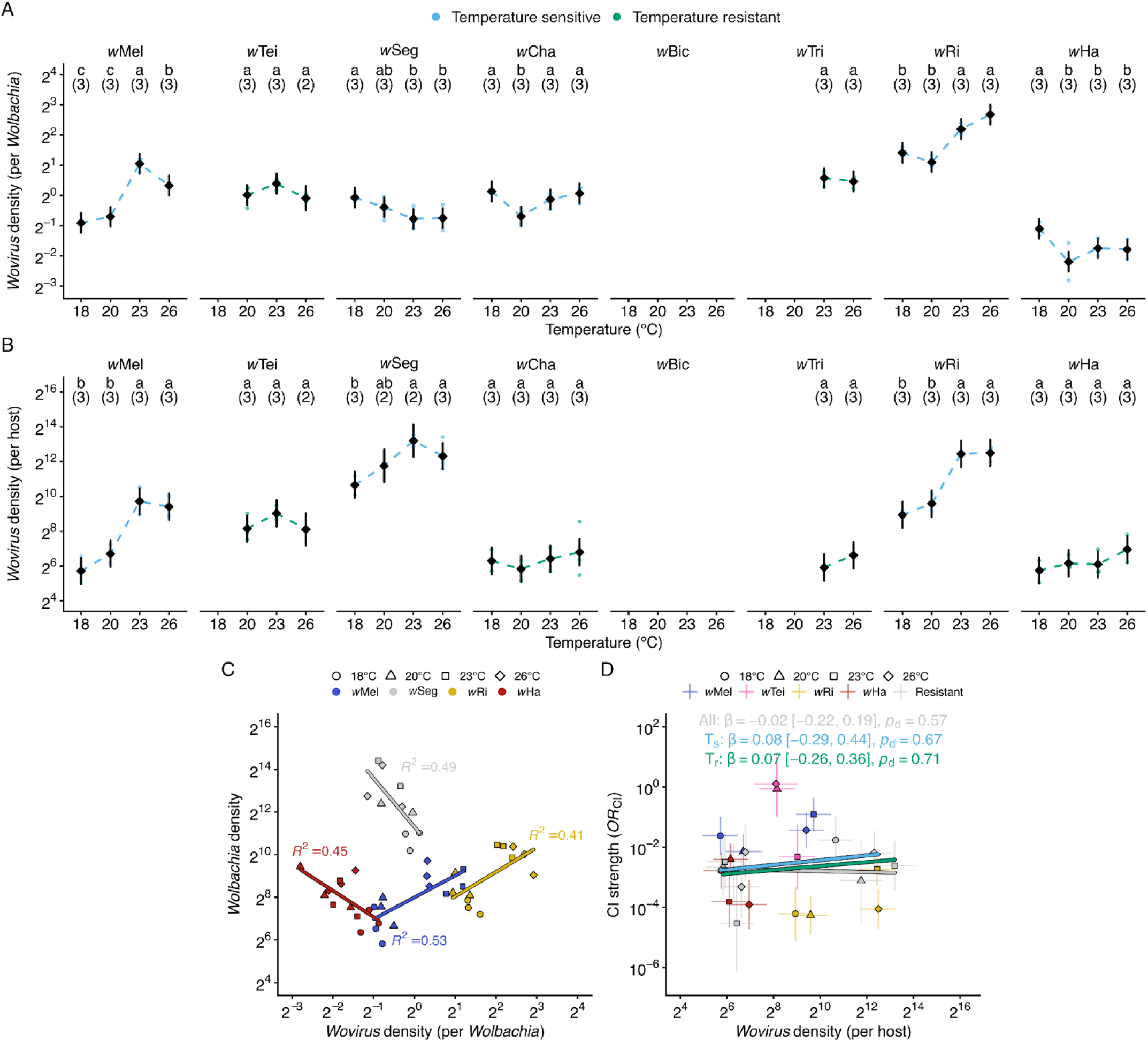
*Wovirus* densities varied with temperature and covaried with *Wolbachia* densities, but did not predict CI strength. **(A)** *Wovirus* densities per *Wolbachia* varied with temperature in five of the seven focal strains (*w*Mel, *w*Seg, *w*Cha, *w*Ri, *w*Ha). **(B)** *Wovirus* densities per host varied with temperature in three of the seven focal strains (*w*Mel, *w*Seg, *w*Ri). **(A, B)** Diamonds represent model-estimated mean densities per strain and temperature, with 95% confidence intervals as vertical lines. Individual points show density values from biological replicates containing 5 pairs of testes. Dashed lines connect estimated means across temperatures to highlight thermal response patterns. Letters indicate significant differences in density among temperatures within each strain (FDR-corrected, α = 0.05); shared letters denote non-significant differences. Sample sizes per group are in parentheses. **(C)** *Wolbachia* densities covaried with *Wovirus* densities per *Wolbachia* across four strains with temperature-sensitive *Wolbachia* densities (*w*Mel, *w*Seg, *w*Ri, *w*Ha). Individual points represent paired *Wolbachia* and *Wovirus* density measurements from the same biological replicates. Regression lines and *R*^2^ values show strain-specific relationships, with all correlations significant (Pearson correlation, *P* < 0.05; *n* = 10 to 12 biological replicates per strain). **(D)** *Wovirus* densities per host did not significantly predict CI strength. Individual points represent estimated marginal means for *OR*_CI_ and *Wovirus* densities per host per strain-temperature combination, with 95% confidence intervals as vertical and horizontal lines. Regression lines and statistics (β [95% credible interval], *p*_d_) were derived from Bayesian phylogenetic mixed models fit separately to all strains (gray; *n* = 25, 7 strains), temperature-sensitive CI strains (T_s;_ light blue; *n* = 15, 4 strains), and temperature-resistant CI strains (T_r;_ green; *n* = 10, 3 strains). **(A-D)** All densities were measured by qPCR. *Wolbachia* densities were calculated as fold-change of *ftsZ* relative to host *nAcRalpha-34E. Wovirus* densities were calculated as fold-change of serine recombinase genes relative to *ftsZ* (per *Wolbachia*) or nAcRalpha-34E (per host). For strains with multiple prophage variants (see Materials & Methods), total *Wovirus* density values were summed across variants (primers: **Table S3**). *w*Bic was excluded because its *Wovirus* serine recombinase gene is incomplete at a contig boundary.

#### Wovirus densities per host varied by strain and thermal exposure

Because the *cif* genes that govern CI are encoded within *Wovirus* prophage genomes, we hypothesized that *Wovirus* copy number per host could influence *cif*-gene dosage and thus CI strength. We therefore also quantified *Wovirus* densities per host (serine recombinase relative to nAcRalpha-34E), which captures total *Wovirus* load in host tissue from both integrated prophage and any extrachromosomal phage genomes. We analyzed how *Wovirus* densities per host vary with strain and temperature using a linear model on log_2_-transformed density values (*N* = 72 samples, each containing 5 pairs of testes; *w*Bic excluded). The model included *Wolbachia* strain (seven focal strains), temperature (18°C, 20°C, 23°C, and 26°C), and their interaction as fixed effects. As with *Wovirus* densities per *Wolbachia*, strain identity was the dominant predictor (ω^2^_p_ = 0.92, *F*_6_ = 137.8, *P* = 4.2e-28), with temperature (ω^2^_p_ = 0.62, *F*_3_ = 39.4, *P* = 7.1e-13) and strain × temperature (ω^2^_p_ = 0.44, *F*_15_ = 4.8, *P* = 1.7e-5) also contributing significantly.

#### Three Wolbachia exhibited temperature-sensitive Wovirus densities per host

Applying the same pairwise comparison framework used for *Wovirus* densities per *Wolbachia*, we determined four of seven strains in this analysis exhibited temperature-resistant per-host *Wovirus* densities, with no significant differences across any temperature comparisons: *w*Tei (ω_2_ = 0.25, largest effect: *RR*_23:26_ = 1.89 [0.67 to 5.27], *P* = 0.20), *w*Cha (ω_2_ = 0, largest effect: *RR*_20:26_ = 0.52 [0.19 to 1.43], *P* = 0.48), *w*Tri (*RR*_23:26_ = 0.62 [0.29 to 1.30], *P* = 0.20), and *w*Ha (ω_2_ = 0.16, largest effect: *RR*_18:26_ = 0.43 [0.16 to 1.20], *P* = 0.17; **Fig 4B**). In contrast, three strains exhibited temperature-sensitive per-host *Wovirus* densities. *w*Mel (ω_2_ = 0.86) showed increased per-host *Wovirus* densities at warmer temperatures, with density at 18°C similar to 20°C (*RR* = 0.50 [0.18 to 1.40], *P* = 0.086) and significantly lower than at 23°C (*RR* = 0.063 [0.023 to 0.17], *P* = 8.9e-9) and 26°C (*RR* = 0.077 [0.028 to 0.21], *P* = 3.4e-8; **Fig 4B**). *w*Ri (ω_2_ = 0.95) exhibited a similar pattern, with density at 18°C similar to 20°C (*RR* = 0.64 [0.23 to 1.77], *P* = 0.28) and significantly lower than at 23°C (*RR* = 0.088 [0.032 to 0.24], *P* = 1.1e-7) and 26°C (*RR* = 0.085 [0.031 to 0.24], *P* = 1.1e-7; **Fig 4B**). In both strains, *Wolbachia* densities and *Wovirus* densities per *Wolbachia* increased at warmer temperatures, producing amplified increases in per-host *Wovirus* densities. *w*Seg (ω_2_ = 0.61) also showed increased per-host *Wovirus* densities at warmer temperatures, with density at 18°C significantly lower than at 23°C (*RR* = 0.17 [0.055 to 0.54], *P* = 6.1e-4) and 26°C (*RR* = 0.32 [0.11 to 0.88], *P* = 0.010; **Fig 4B**). This is notable because *w*Seg *Wovirus* densities per *Wolbachia* decreased at warmer temperatures. Overall, the three strains with temperature-sensitive per-host *Wovirus* densities (*w*Mel, *w*Seg, and *w*Ri) showed higher total *Wovirus* load at warmer temperatures. Temperature-sensitive and temperature-resistant classes include both *w*Mel-like and *w*Ri-like *Wolbachia*.

#### Temperature-sensitive Wolbachia densities covaried with Wovirus densities

To test whether *Wovirus* dynamics explain temperature-sensitive *Wolbachia* density variation, we used Bayesian phylogenetic mixed models to test whether *Wovirus* densities per *Wolbachia* predict *Wolbachia* densities across strains. We focused on six strains where both measurements were available (*w*Mel, *w*Seg, *w*Cha, *w*Ri, *w*Ha, and *w*Tei), excluding *w*Bic (no *Wovirus* measurements) and *w*Tri (only two temperature points). Bayesian analysis revealed no consistent overall relationship between *Wovirus* densities per *Wolbachia* and *Wolbachia* densities when considering all strains (β = 0.02 [−0.39 to 0.43], *R*^2^ = 0.85, *p*_d_ = 0.54, *n* = 66, 6 strains) or strains with temperature-sensitive *Wolbachia* densities (β = −0.10 [−0.51 to 0.31], *R*^2^ = 0.88, *p*_d_ = 0.69, *n* = 46, 4 strains). Credible intervals spanned zero in both cases, and *p*_d_ values indicated weak directional certainty, providing no support for a generalizable *Wovirus*-*Wolbachia* relationship across strains.

To complement cross-strain analyses, we examined within-strain relationships using Pearson correlations on raw qPCR data (*n* = 8 to 12 per strain). Within-strain correlations showed distinct patterns. Two strains exhibited positive covariation, where higher *Wovirus* densities corresponded to higher *Wolbachia* densities: *w*Mel (*r* = 0.73, *R*^2^ = 0.53, *P* = 0.007, *n* = 12) and *w*Ri (*r* = 0.64, *R*^2^ = 0.41, *P* = 0.025, *n* = 12; **Fig 4C**). In contrast, two strains showed negative covariation, where higher *Wovirus* densities correspond to lower *Wolbachia* densities: *w*Seg (*r* = −0.70, *R*^2^ = 0.49, *P* = 0.023, *n* = 10) and *w*Ha (*r* = −0.67, *R*^2^ = 0.45, *P* = 0.017, *n* = 12; **Fig 4C**). Finally, two strains showed no significant relationship: *w*Cha (*r* = 0.08, *R*^2^ = 0.01, *P* = 0.81, *n* = 12) and *w*Tei (*r* = −0.30, *R*^2^ = 0.09, *P* = 0.46, *n* = 8). The opposing directions across strains help explain why no overall relationship emerged in the Bayesian models. These results suggest that temperature-sensitive *Wovirus*-*Wolbachia* dynamics differ among *Wolbachia* strains, with some showing positive and others negative covariation, including closely related *w*Mel and *w*Seg that show distinct patterns.

#### Wovirus densities per host did not predict CI strength

Given that *cif* genes are encoded within *Wovirus* genomes, we tested whether *Wovirus* densities per host predict CI strength using the same Bayesian phylogenetic mixed model and within-strain concordance frameworks described above. Bayesian analysis revealed no relationship between *OR*_CI_ and *Wovirus* densities per host when considering all strains (β = −0.02 [−0.22 to 0.19], *R*^2^ = 0.47, *p*_d_ = 0.57, *n* = 25, 7 strains), temperature-sensitive CI strains (β = 0.08 [−0.29 to 0.44], *R*^2^ = 0.63, *p*_d_ = 0.67, *n* = 15, 4 strains), or temperature-resistant CI strains (β = 0.07 [−0.26 to 0.36], *R*^2^ = 0.20, *p*_d_ = 0.71, *n* = 10, 3 strains; **Fig 4D**). All credible intervals included zero, and *p*_d_ values indicated little confidence in the direction of effects. Within-strain concordance analysis similarly showed inconsistent relationships between *Wovirus* densities per host and CI strength, with no strain exhibiting perfect concordance. Only *w*Mel showed significant concordance (τ = +0.62, *p*_d_ = 0.98), but in the positive direction, where higher *Wovirus* densities per host correspond to weaker CI and opposite to the prediction that more *Wovirus* copies increase *cif* dosage and strengthen CI. The remaining five strains with *Wovirus* data showed no significant concordance: *w*Ri (τ = +0.63, *p*_d_ = 0.94), *w*Tei (τ = −0.95, *p*_d_ = 0.94), *w*Seg (τ = −0.46, *p*_d_ = 0.67), *w*Ha (τ = −0.51, *p*_d_ = 0.67), and *w*Cha (τ = +0.12, *p*_d_ = 0.73). Thus, despite *Wovirus* genomes encoding the *cif* genes responsible for CI, total *Wovirus* load per host does not predict CI-strength variation across temperatures.

### Temperature modulated *cifB* transcript levels

#### cifB transcript abundance in temperature-sensitive CI strains varied by gene variant and temperature

The *cifB* gene encodes the toxin responsible for CI-induced embryonic lethality (Beckmann et al. 2017; LePage et al. 2017; Shropshire and Bordenstein 2019; Adams et al. 2021; Sun et al. 2022). We hypothesized that temperature-sensitive effects on *cifB* transcription could have influenced the temperature-sensitive CI-strength variation we observed. Before testing this hypothesis, we characterized *cifB* transcription variation from the three temperature-sensitive CI *Wolbachia* strains that induced CI at all tested temperatures (*w*Mel, *w*Ri, and *w*Ha). We dissected testes from male siblings of flies used in our CI experiments, extracted total RNA, and quantified *cifB*-transcript abundance relative to a synthetic RNA spike-in control via RT-ddPCR. For each strain, we compared transcript levels at two temperatures: 23°C versus either 18°C (*w*Mel and *w*Ri) or 20°C (*w*Ha). We analyzed *cifB*-transcript levels across temperature using a linear model on log_2_-transformed transcript values (*N* = 95 *cifB* measurements). The model included *cifB* variant (five variants: *cifB*_*w*Mel[T1]_, *cifB*_*w*Ri[T1]_, *cifB*_*w*Ri[T2]_, *cifB*_*w*Ha[T1-1]_, and *cifB*_*w*Ha[T1-2]_), temperature treatment (cool vs warm), and their interaction as fixed effects. In order of decreasing effect size (ω^2^_p_), significant terms included: *cifB* variant (ω^2^ = 0.90, *F*_4_ = 225.4, *P* = 2.2e-44), temperature (ω^2^_p_ = 0.49, *F*_1_ = 91.6, *P* = 3.8e-15), and *cifB* variant × temperature (ω^2^_p_ = 0.23, *F*_4_ = 7.9, *P* = 1.7e-5). Hence, genetic variation significantly predicts *cifB*-transcript abundance, and temperature affects *cifB* dosage in an variant-specific manner.

#### Four of the five cifB variants exhibited temperature-sensitive transcript abundance

Having established that temperature affected *cifB* transcription overall, we next examined which specific variants showed temperature-sensitive transcript variation by comparing estimated marginal means between cool and warm temperatures (*RR*; > 1 = higher transcription at warm, < 1 = higher transcription at cool). Of the five *cifB* variants measured, four exhibited temperature-sensitive transcription: *cifB*_*w*Mel[T1]_ (*RR* = 0.45 [0.29 to 0.70], *P* = 5.9e-4), *cifB*_*w*Ri[T1]_ (*RR* = 0.18 [0.11 to 0.28], *P* = 1.1e-10), *cifB*_*w*Ri[T2]_ (*RR* = 0.27 [0.17 to 0.43], *P* = 4.3e-7), and *cifB*_*w*Ha[T1-1]_ (*RR* = 0.32 [0.21 to 0.49], *P* = 1.6e-6) were each more highly expressed at the cooler temperature (18°C for *w*Mel and *w*Ri, 20°C for *w*Ha) compared to the warmer temperature (23°C for all strains; **Fig 5A**). In contrast, *cifB*_*w*Ha[T1-2]_ transcript levels were similar between 20°C and 23°C (*RR* = 0.98 [0.63 to 1.51], *P* = 0.91). Overall, four of five *cifB* variants exhibited temperature-sensitive transcript abundance, with elevated transcription at cooler temperatures.

**Figure 5.**
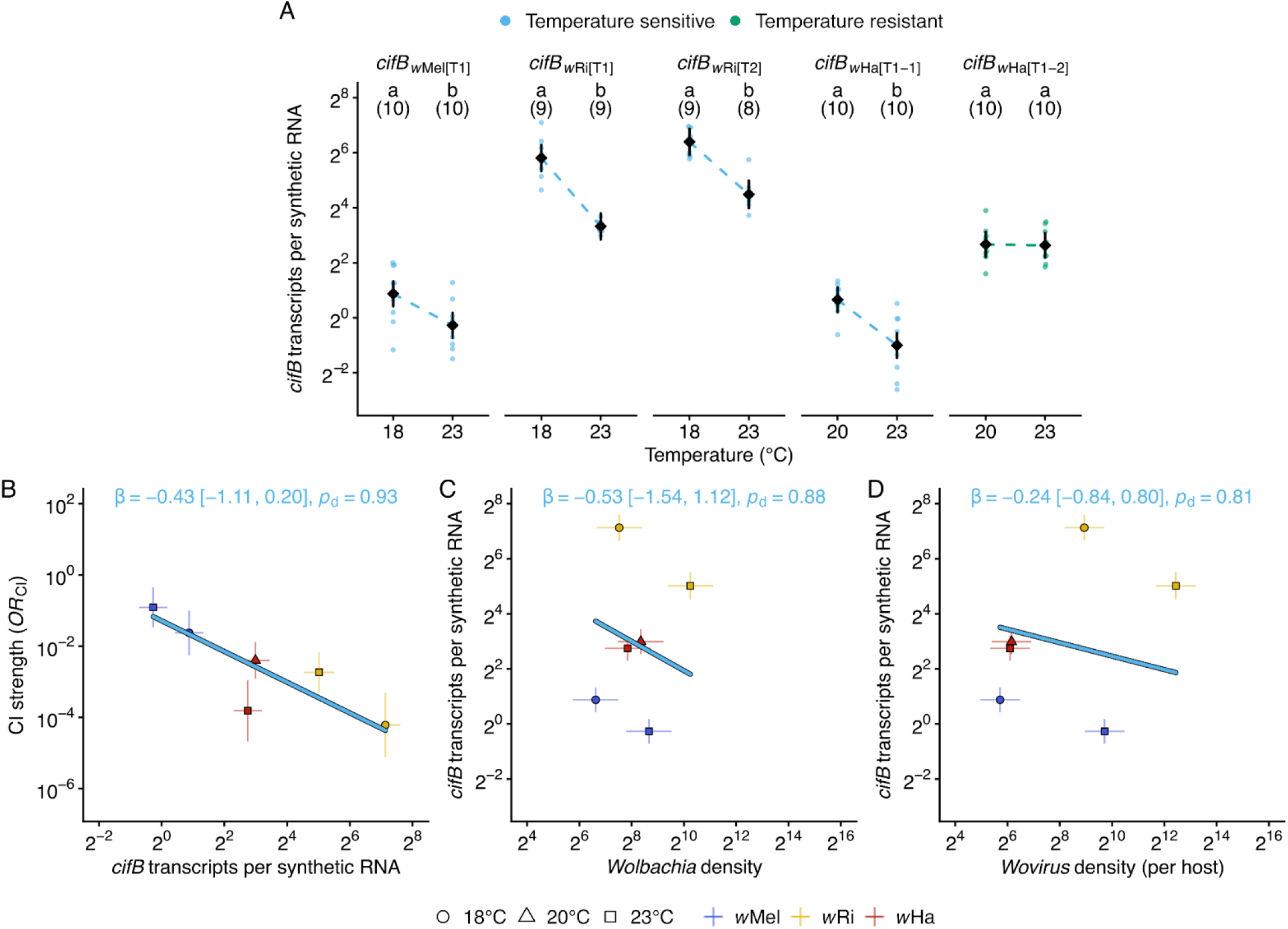
*cifB*-transcript levels varied with temperature, partly predicted CI strength, and were decoupled from *Wolbachia* and *Wovirus* densities. **(A)** *cifB*-transcript abundance varied with temperature across five *cifB*-gene variants (see Materials & Methods) from three temperature-sensitive CI strains that exhibited CI at more than one temperature (*w*Mel, *w*Ri, *w*Ha). Diamonds represent model-estimated mean transcript abundances per variant and temperature, with 95% confidence intervals as vertical lines. Individual points show transcript measurements from biological replicates. For *w*Mel and *w*Ri, comparisons were made between 18°C and 23°C; for *w*Ha, comparisons were made between 20°C and 23°C. Dashed lines connect estimated means across temperatures. Letters indicate significant differences in transcript abundance among temperatures within each variant (FDR-corrected, α = 0.05); shared letters denote non-significant differences. Sample sizes per group are in parentheses. **(B)** Higher *cifB*-transcript abundance partly predicted stronger CI (lower *OR*_CI_) across temperature-sensitive CI strains. **(C)** *cifB*-transcript abundance was not significantly associated with *Wolbachia* densities. **(D)** *cifB*-transcript abundance was not significantly associated with *Wovirus* densities per host. **(B-D)** Individual points represent estimated marginal means per strain-temperature combination, with 95% confidence intervals as vertical and horizontal lines. Regression lines and statistics (β [95% credible interval], *p*_d_) are from Bayesian phylogenetic mixed models. **(A-D)** *cifB*-transcript abundance was quantified by RT-ddPCR, normalized to a synthetic RNA spike-in control. *Wolbachia* and *Wovirus* densities were measured by qPCR as fold-change of *ftsZ* and serine recombinase genes, respectively, relative to host nAcRalpha-34E (primers: **Table S3**).

#### cifB transcription partly predicted CI-strength variation in strains with temperature-sensitive CI

These results established that *cifB* transcription varies with temperature in most variants we analyzed. We next tested whether this variation predicted CI strength using Bayesian phylogenetic mixed models to analyze three temperature-sensitive CI strains: *w*Mel, *w*Ri, and *w*Ha. Bayesian analysis reveals a negative relationship between *cifB* transcription and OR_CI_ (*β* = −0.43 [−1.11 to 0.20], *R*^2^ = 0.66, *p*_d_ = 0.93, *n* = 6, 3 strains; **Fig 5B**), indicating that higher *cifB* transcript abundance was associated with stronger CI (lower *OR*_CI_). Although the 95% credible interval included zero, the probability of direction indicated 93% posterior confidence that the true effect was negative. Within-strain concordance analysis using Kendall’s τ with Monte Carlo propagation of uncertainty supported this pattern for two of three strains: *w*Mel (τ = −1.00, *p*_d_ = 0.99, *n* = 2) and *w*Ri (τ = −1.00, *p*_d_ = 1.00, *n* = 2) both showed complete negative concordance between *cifB* transcription and CI strength, where higher *cifB* at cooler temperatures corresponded to stronger CI. In contrast, *w*Ha showed positive discordance (τ = +1.00, *p*_d_ = 0.99, *n* = 2), where higher *cifB* corresponded to weaker CI. These results indicate that *cifB* transcription partly predicts temperature-sensitive CI-strength variation.

#### Neither Wolbachia nor Wovirus per host densities predicted cifB transcription

If *cifB* transcription predicts CI-strength variation but *Wolbachia* densities do not, this suggests a disconnect between bacterial abundance and *cifB*-transcript levels across temperatures. Applying the same Bayesian and concordance frameworks with *cifB* transcription as the response, we did not find a positive relationship between *cifB* transcription and *Wolbachia* densities (β = −0.53 [−1.51 to 1.12], *R*^*2*^ = 0.86, *p*_d_ = 0.88, *n* = 6, 3 strains; **Fig 5C**). The probability of a positive effect was only 12%. Concordance analysis revealed significant negative concordance for *w*Mel (τ = −1, *p*_d_ = 1.00) and *w*Ri (τ = −1, *p*_d_ = 1.00), where higher *Wolbachia* densities corresponded to lower *cifB* transcription, while *w*Ha showed non-significant negative concordance (τ = −0.83, *p*_d_ = 0.66). Thus, *cifB*-transcript levels did not increase with *Wolbachia* densities and instead declined significantly in 2 of 3 strains, potentially explaining why bacterial abundance failed to predict CI strength. Because *cif* genes are encoded within *Wovirus* prophage genomes, we further asked whether total *Wovirus* densities per host predicted *cifB* transcription. Neither the Bayesian model (*β* = −0.24 [−0.84 to 0.80], *R*^2^ = 0.81, *p*_d_ = 0.81, *n* = 6, 3 strains; **Fig 5D**) nor the concordance analysis supported such a relationship, with the credible interval spanning zero symmetrically; and while *w*Mel (τ = −1.00, *p*_d_ = 1.00) and *w*Ri (τ = −1.00, *p*_d_ = 1.00) again showed perfect negative concordance, *w*Ha did not (τ = −0.71, *p*_d_ = 0.62). Total phage load is therefore decoupled from the expression of phage-encoded *cif* genes, mirroring the pattern observed between *Wolbachia* density and *cifB* transcription.

## Discussion

Temperature affects critical biochemical and cellular processes in ecotherms (*e.g*., Krogh 1916; Clarke and Fraser 2004; Cooper et al. 2012; Somero et al. 2017), making it an important factor for the physiology and fitness of the *Drosophila* hosts in our study (Klepsatel et al. 2013; Tobler et al. 2015; Klepsatel et al. 2016). Similar to interactions between nuclear and microbe-derived mitochondrial genomes (Hoekstra et al. 2013), temperature is an important modulator of endosymbiont-host interactions and phenotypes such as CI (Clancy and Hoffmann 1998; Bordenstein and Bordenstein 2011; Ross et al. 2017; Nasehi et al. 2022), yet the underlying mechanisms remain poorly understood. Our findings across eight *Wolbachia*-*Drosophila* systems spanning ~7.5 million years of *Wolbachia* divergence indicate that temperature effects on CI are system-dependent, with closely related variants often differing in their responses. Understanding this variation will likely assist efforts to interpret variation in *Wolbachia* frequencies through space and time (*e.g*., Kriesner et al. 2013; Kriesner et al. 2016; Schuler et al. 2016; Cooper et al. 2017; Ross et al. 2019b). Among candidate predictors of variable CI strength, neither development times nor *Wolbachia* densities reliably predicted CI strength, despite both varying significantly with temperature. Instead, *cifB* transcription partly explained temperature-sensitive CI. Notably, *cifB*-transcript levels were decoupled from *Wolbachia* densities, and *Wovirus* densities covaried with *Wolbachia* in strain-specific ways. We discuss these findings, and others, below.

### System-dependent temperature effects on the strength of CI and CI rescue

#### Temperature influences the egg hatch of compatible crosses

Evaluating contributions of CI to the spread and maintenance of *Wolbachia* first requires establishing baseline embryonic survival. Variation in compatible egg hatch across temperatures likely reflects species-specific thermal performance, where fertility limits are often narrower than survival limits (van Heerwaarden and Sgrò 2021; Klepsatel et al. 2023). Our observation of thermally stable egg hatch for compatible *D. melanogaster* crosses is consistent with prior observations of the relatively high fecundity following development between 22°C and 26°C (Cooper et al. 2010; Klepsatel et al. 2023). While effects were small, the two *D. simulans* genotypes differed in their responses, with the *w*Ha-genotype showing stable hatching across temperatures and the *w*Ri-genotype showing reduced hatching at 18°C, consistent with documented population differences in hatchability (Austin and Moehring 2013). These *D. simulans* genotypes also carry distinct mitochondrial haplotypes (siI in the *w*Ha-genotype, siII in the *w*Ri-genotype; Ballard 2004) that are known to differ in thermal physiology (Pichaud et al. 2010) and organismal fitness traits (James and Ballard 2003; Ballard et al. 2007), making it difficult to attribute the observed difference solely to nuclear background. Among the five species without prior thermal hatching data, *D. teissieri* exhibited the most dramatic sensitivity. Hatching at 23°C was robust but declined at 20°C and at 26°C, where very few eggs hatched. This narrow optimum aligns with *D. teissieri*’s ecology as a forest-restricted specialist, with reduced fecundity and egg-to-adult viability observed at temperatures above 24°C (Cooper et al. 2018). *D. bicornuta* from Taiwan also displayed reductions in egg hatch at 26°C, though less severe, while lower *D. seguyi* compatible hatch at 18°C was statistically nonsignificant. In contrast, *D. chauvacae* from Madagascar and *D. triauraria* from Japan both maintained stable hatching across tested temperatures. However, failure of *D. triauraria* to develop at 18°C and 20°C suggests this species is highly sensitive to cool temperatures. Collectively, the egg hatch of compatible crosses varies substantially among host systems and temperatures, independent of *Wolbachia* effects.

#### Temperature influences the strength of CI in four systems

After accounting for baseline variation, four strains exhibited temperature-sensitive CI (*w*Mel, *w*Tei, *w*Ri, *w*Ha). *w*Tei exhibited the most thermally restricted CI, detectable only at 23°C. However, reduced compatible-cross hatching at 20°C and 26°C limits statistical power to detect CI at those temperatures, inhibiting us from distinguishing true absence of CI from an inability to detect it. This suggests that temperature — in addition to *D. teissieri* nuclear effects (Cooper et al. 2017) — contributes to differences in the detection of CI across studies (Zabalou et al. 2004; Cooper et al. 2017), underscoring the necessity of testing for CI across contexts. In contrast, closely related *w*Mel caused CI at all temperatures, but CI strength was lower at 23°C than at 20°C. CI induced by *w*Mel transinfections in *Ae. aegypti* are reduced at hot temperatures (Ross et al. 2017), and future work focused on effects of temperature stress on CI strength in *w*Mel-*D. melanogaster* and other natural systems are needed. The *w*Ri-*D. simulans* genotype caused weaker CI at 23°C than at the other three temperatures. Prior *w*Ri studies reported reduced CI strength at 27°C (Clancy and Hoffmann 1998) and 28°C (Hoffmann et al. 1986), suggesting CI strength is likely to be lower at hotter temperatures than we tested. In contrast, *w*Ha tended to show the weakest CI at 20°C, with stronger CI induced at both 23°C and 26°C. This finding agrees with prior analyses focused on single temperatures (23°C and 25°C) that found strong CI (O’Neill and Karr 1990; Martinez et al. 2015; Shropshire et al. 2022). Among the four temperature-resistant strains (*w*Seg, *w*Cha, *w*Bic, *w*Tri), *w*Cha and *w*Bic have not been tested for CI at any temperature, but both strains were predicted to cause CI based on their *cif* profiles (Martinez et al. 2021; Shropshire et al. 2026). We established that both cause relatively strong and stable CI across temperatures. Similar to estimates at 25°C (Shropshire et al. 2026), *w*Seg caused strong and stable CI across all four temperatures. Finally, *w*Tri caused relatively strong CI at 23°C and 26°C, where tests were possible. We conclude that temperature often modulates CI strength, but the specific responses vary widely among strains. This includes closely related *w*Mel-like (*i.e., w*Mel and *w*Tei versus *w*Seg and *w*Cha) and very closely related *w*Ri-like variants (*i.e., w*Ri versus *w*Tri) that may cause temperature-sensitive or temperature-resistant CI.

#### Temperature influences CI-rescue efficiency in four systems

CI is only advantageous to *Wolbachia* if symbiotic females can rescue it (Hoffmann et al. 1990). Studies commonly demonstrate complete rescue, even when CI strength varies (*e.g*., Clancy and Hoffmann 1998; Shropshire et al. 2021a). Four strains in our study exhibited complete rescue across all tested temperatures (*w*Mel, *w*Tei, *w*Seg, *w*Tri). However, four show incomplete rescue at one or more temperatures: *w*Bic and *w*Ha at 23°C, *w*Cha at 20°C, and *w*Ri at both 20°C and 26°C. Our *w*Ri findings contrast with those of Clancy and Hoffmann (1998), who did not observe temperature effects on rescue. Their design reared both sexes at the same temperature, while we reared males at experimental temperatures (18°C to 26°C) and all females at 23°C. Because temperature alters *cif* transcript levels, we hypothesize that rearing sexes at different temperatures creates a mismatch between CifB in sperm and CifA in the embryo, resulting in incomplete rescue. However, temperature differences between sexes cannot explain incomplete rescue by *w*Bic and *w*Ha-carrying females, where both sexes were reared at 23°C. Compared to variation in CI strength, incomplete rescue is rare but documented in other systems: field heat stress in transinfected *Ae. aegypti* (Ross et al. 2019a), temperature extremes in *Encarsia* wasps with *Cardinium* (Doremus et al. 2019), and transinfected *w*Tei in a novel *D. simulans* background (Zabalou et al. 2008). *cif*-dosage mismatches may still arise when conditions produce sex-specific differences in symbiont physiology or host interactions, potentially explaining incomplete rescue by *w*Bic and *w*Ha in host females. These findings demonstrate that temperature effects extend beyond CI induction to rescue modulation. Accounting for such effects when modeling *Wolbachia* dynamics (*e.g*., Hoffmann et al. 1990) may improve our ability to understand the factors governing variable *Wolbachia* frequencies observed in many systems (Kriesner et al. 2013; Schuler et al. 2016; Cooper et al. 2017; Wheeler et al. 2021; Hague et al. 2022; Sanaei et al. 2022; Shastry et al. 2022).

### Mechanisms underlying temperature-sensitive CI strength

#### Development time varies with temperature and Wolbachia but does not predict CI

We first examined development time as a candidate predictor (Doremus et al. 2019; Doremus et al. 2020; Shropshire et al. 2022). In all systems, development time covaried with temperature, yet two *Wolbachia* strains unexpectedly accelerated host development at specific temperatures: *w*Cha shortened development by 1.9 days at 20°C and 1.3 days at 26°C, while *w*Bic accelerated development by 3.6 days at 18°C only. The remaining four systems (*w*Mel, *w*Tei, *w*Seg, *w*Ri) showed no detectable effect, consistent with prior reports in *w*Mel (Harcombe and Hoffmann 2004; Strunov et al. 2022). Both effects represented acceleration rather than delay, though *Wolbachia* effects on development time vary in direction across systems, with some strains accelerating (Nikoh et al. 2014; Hickin et al. 2022; Lindsey et al. 2025) and others delaying development (Reynolds et al. 2003; Cao et al. 2019). Possible mechanisms include nutrient provisioning and host metabolic reprogramming (Ponton et al. 2015; Newton and Rice 2020; Lindsey et al. 2025; Niu et al. 2025). Notably, *w*Cha and *w*Bic effects appeared at our thermal limits, suggesting that *Wolbachia* may buffer host development against thermal stress, as documented for host survival in other strains (Gruntenko et al. 2017; Burdina et al. 2021). Collectively, these results indicate *Wolbachia* can provide developmental benefits under thermal stress in a strain-specific manner.

While development time correlates with temperature, it did not predict CI strength in our dataset, either across or within strains. However, other studies have demonstrated such correlations: longer pupal development yielded stronger CI in *Cardinium*-bearing *Encarsia* wasps, whether achieved through temperature or juvenile hormone treatment (Doremus et al. 2019; Doremus et al. 2020); longer larval development weakly correlated with stronger CI across divergent *Wolbachia*-*Drosophila* strains (Shropshire et al. 2022); and cold-induced prolongation of development increased CI despite reducing *Wolbachia* densities in *Nasonia* wasps (Bordenstein and Bordenstein 2011). Yet Yamada et al. (2007) found the opposite pattern: stronger CI in faster-developing *D. melanogaster* males, but no effect in *D. simulans*. This suggests that development time may covary with unmeasured factors that vary among systems and contexts. CI induction is associated with potentially time-restricted processes during spermatogenesis, including *Wolbachia* proliferation within developing spermatocyte cysts (Clark et al. 2002; Clark et al. 2003) and Cif modification of sperm chromatin and loading into maturing sperm (Xiao et al. 2021; Kaur et al. 2022; Terretaz et al. 2023; Kaur et al. 2024). Temperature and other factors that modulate development time may differentially affect the duration of these spermatogenic windows, altering the time available for Cif accumulation and/or action.

#### Wolbachia and Wovirus densities vary with temperature but do not predict CI strength

Beyond the relationship with development time described above, we evaluated the hypothesis that more *Wolbachia* in testes yields stronger CI. Focal *Wolbachia* exhibited diverse density responses to temperature: cold inhibition (*w*Mel, *w*Seg, *w*Ri, *w*Ha), intermediate peak (*w*Bic), and temperature-resistance (*w*Tei, *w*Cha, *w*Tri). Temperature-density relationships have been observed for *w*Mel, where density declined at higher temperatures in transinfected *Ae. aegypti* (Ross et al. 2017), and for *w*Ri, where density varied with rearing temperature in *D. simulans* embryos (Clancy and Hoffmann 1998). The causes of temperature-sensitive *Wolbachia* densities are unknown, and we hypothesized that *Wovirus* activity could plausibly contribute (Bordenstein and Bordenstein 2022). *Wovirus* densities varied with temperature in five of seven focal strains (*w*Mel, *w*Seg, *w*Cha, *w*Ri, *w*Ha). Although qPCR cannot distinguish phage life-cycle states, the direction of *Wovirus*-*Wolbachia* correlations is informative. In *w*Seg and *w*Ha, *Wovirus* density per *Wolbachia* increased as *Wolbachia* densities decreased. These negative correlations are consistent with temperature-triggered lytic activity, as documented in *Wolbachia* of *N. vitripennis* (Bordenstein and Bordenstein 2011), *H. hebetor* (Nasehi et al. 2022), and *T. urticae* (Lu et al. 2012). In contrast, in *w*Mel and *w*Ri, *Wovirus* density per *Wolbachia* increases with *w*Mel and *w*Ri density, consistent with non-lytic episomal replication (Biliske et al. 2011; Mäntynen et al. 2021).

However, *Wolbachia* and phage replication could also respond to temperature independently; indeed, *w*Cha exhibited temperature-sensitive *Wovirus* density despite stable *Wolbachia* density. More broadly, temperature-sensitive *Wolbachia* densities likely reflect both bacterial and host determinants. On the bacterial side, beyond *Wovirus* dynamics, variation in Octomom copy number directly modulates *Wolbachia* replication rate and densities (Chrostek and Teixeira 2015); while on the host side, autophagy-mediated regulation (Voronin et al. 2012; Deehan et al. 2021) and the maternal effect gene *Wds*, which dominantly suppresses *Wolbachia* densities in *Nasonia* (Funkhouser-Jones et al. 2018), may plausibly contribute to the variation. Notably, CifB modulates host autophagy (Deehan et al. 2021), suggesting *Wolbachia* genotypes can feed back on host pathways that control *Wolbachia* densities. The strain-specific thermal density profiles observed here likely reflect differential *Wolbachia* replication dynamics, bacterial genetic architecture, and host regulatory mechanisms, each of which could respond to temperature.

Regardless of the mechanisms underlying temperature-dependent density modulation, our goal was to evaluate the *Wolbachia* density hypothesis (Breeuwer and Werren 1993). While *Wolbachia* densities have covaried with CI in diverse contexts (*e.g*., Clancy and Hoffmann 1998; Yamada et al. 2007; Bordenstein and Bordenstein 2011; Layton et al. 2019), we found no instances where higher densities consistently corresponded to stronger CI. We observed the strongest associations in *w*Mel and *w*Ri, but in the direction opposing the density hypothesis: higher densities corresponded to weaker CI, paralleling the observation that *w*Mel testes’ density increased as CI strength declined with *D. melanogaster* male age (Shropshire et al. 2021a). Density-CI decoupling has been documented across diverse experimental contexts: two-fold temperature-induced density differences did not correspond with changes in CI strength in *Leptopilina* (Mouton et al. 2006); both cool and warm temperatures reduced *Cardinium* density in *E. suzannae*, yet had opposing effects on CI strength (Doremus et al. 2019); experimentally reducing *Wolbachia* density via larval crowding did not weaken CI in *Ae. albopictus* (Dutton and Sinkins 2004); and CI in *D. melanogaster* varied from 95% to undetectable across development times, despite no density or localization differences in testes (Yamada et al. 2007). This last observation is particularly relevant because it demonstrates that even *Wolbachia* distribution within spermatocyte cysts (Clark et al. 2002; Clark et al. 2003) fails to explain CI-strength variation in a development-time context, though temperature-dependent shifts in localization relative to Cif-protein action during sperm chromatin remodeling and loading (Xiao et al. 2021; Kaur et al. 2022; Terretaz et al. 2023) remain untested. We conclude that *Wolbachia* and *Wovirus* densities alone are insufficient to predict CI-strength variation across temperatures.

### cifB transcription partly predicts CI variation and is decoupled from Wolbachia density

With neither development time nor *Wolbachia* density predicting CI, we tested the *cifB*-dosage hypothesis that higher *cifB*-transcript abundance in testes yields stronger CI. We observed temperature-sensitive *cifB* transcription in all strains that we tested: four of five gene variants showed transcript declines at warmer temperatures, with only *cifB*_*w*Ha[T1-2]_ remaining relatively stable. While the overall cross-strain relationship between *cifB* dosage and CI strength remains uncertain (93% probability of direction), clear patterns emerged at the strain level. For *w*Mel and *w*Ri, temperature-sensitive *cifB*-transcript levels were concordant with CI strength, with higher *cifB* abundance at cooler temperatures corresponding to stronger CI. In contrast, for *w*Ha the net *cifB* dosage difference between temperatures with statistically different CI strengths was modest and confidence intervals overlapped. These data align with prior evidence that *cifB*-transcript levels better predict CI strength than do *Wolbachia* densities, though as with our data, unaccounted for variation in CI-strength remains. *cifB* dosage predicts CI strength across strains under fixed rearing conditions, but exceptions exist (Shropshire et al. 2022). For example, *w*Bai of *D. baimaii* induces very strong CI despite *cifB*-transcript levels significantly lower than those of weak-CI-inducing *w*Mel (Shropshire et al. 2022). High *cifB* transcription also corresponded to strong CI as male *D. simulans* carrying *w*Ri aged (Shropshire et al. 2021a), but *cifB* transcription failed to predict age-dependent CI-strength variation induced by *w*Mel in *D. melanogaster* and *Wolbachia* in *H. hebetor* (Shropshire et al. 2021a; Nasehi et al. 2022).

Cross-strain variation in the dosage-CI relationship likely reflects *cifB* genetic architecture, as CifB enzymatic activity and sequence divergence drive variation in transgenic CI strength among homologs (Shropshire et al. 2020; Beckmann et al. 2021; Shropshire et al. 2021b; Shropshire et al. 2022). Both cross-strain and within-strain divergences from the dosage model may also reflect spatiotemporal *cif* expression patterns during spermatogenesis (Shropshire and Bordenstein 2019; Xiao et al. 2021; Kaur et al. 2022; Terretaz et al. 2023; Kaur et al. 2024), which testes-level measurements cannot resolve. That testes-level *cifB* dosage in *w*Mel explains temperature-sensitive CI but not male-age-dependent CI underscores the model’s context-dependence, suggesting that testes-level *cifB* transcription in complex field environments is likely insufficient to predict CI strength across all contexts. *cifB* dosage remains the strongest predictor of CI strength identified to date, yet gaps between dosage and phenotype point to a hierarchical model in which *cifB* dosage sets CI potential while protein activity and localization modulate the realized phenotype.

Since neither bacterial nor phage density accounted for *cifB*-transcript variation, regulation likely operates directly on transcription. Despite their reduced genomes, *Wolbachia* retain environmentally responsive transcriptional regulation, including the heat shock sigma factor RpoH (Lindsey 2020). The *cifA*-*cifB* gene pair is organized as a bicistronic operon within *Woviruses*, and a predicted Rho-independent transcription terminator in the intergenic region may attenuate read-through to *cifB*, enabling differential regulation of the two genes, consistent with dosage differences between *cifA* and *cifB* observed across host development (Gutzwiller et al. 2015; Lindsey et al. 2018). Lindsey (2020) hypothesized that this terminator is actively modulated, as *Wolbachia* encode Nus transcription elongation factors (NusA, NusB, NusG) that could toggle between termination after *cifA* and antitermination allowing full *cifA*-*cifB* read-through. Suliman et al. (2025) confirmed this architecture via transcription start-site mapping in *w*MelPop-CLA and *w*AlbB, identifying a single start site upstream of *cifA* with no independent *cifB* promoter in either strain. The *cif*-operon start site in *w*AlbB was downregulated at 34°C, demonstrating temperature-sensitive regulation of *cif*-transcription initiation (Suliman et al. 2025). While *Wolbachia* and *Wovirus* abundance may impact *cifB*-transcript loads in some systems and contexts, we hypothesize that *cifB* regulation depends on environmental stimuli that govern transcriptional initiation and termination.

## Conclusions

Our results demonstrate that temperature shapes *Wolbachia*-induced CI through strain-specific effects, with *cifB*-transcript levels explaining part of this variation. Downstream factors — including protein activity and subcellular localization — likely further modulate the realized phenotype. Together with our observations of temperature-sensitive rescue, *Wolbachia*-associated developmental acceleration at thermal extremes, strain-specific *Wovirus*-*Wolbachia* covariance patterns, and decoupling of *cifB* transcription from *Wolbachia*/*Wovirus* density, these findings reveal temperature as a pervasive modulator of *Wolbachia*-host interactions.

These results motivate research in several directions. First, while we document system-specific variation at multiple levels, studies that assess these and additional traits across broader *Wolbachia*-host combinations, wider temperature ranges, and reciprocally introgressed host genotypes will be important for assessing generality. Because mitochondrial haplotypes that co-transmit with *Wolbachia* can independently affect host thermal physiology (James and Ballard 2003; Pichaud et al. 2010), microinjection experiments that decouple these cytoplasmic components will be needed to disentangle their respective contributions. Host background effects on CI are already well documented (*e.g*., Reynolds and Hoffmann 2002; Cooper et al. 2017; Wybouw et al. 2022), with the success of *w*Mel-based biocontrol depending on strong CI expressed in transinfected *Ae. aegypti* backgrounds (Walker et al. 2011; Utarini et al. 2021; de Morais Batista et al. 2026). Second, the thermal sensitivity we observe suggests that considering *Wolbachia* (and other endosymbionts) when analyzing arthropod responses to temperature may reveal endosymbiont contributions to trait variation. Third, explaining *Wolbachia* frequencies, which often fluctuate through time and space, will benefit from incorporating temperature effects on the traits that govern them. Finally, while controlled laboratory experiments facilitate mechanistic dissection of these interactions, they are unlikely to produce results that fully reflect dynamics under natural conditions. The patterns we observed — combined with growing evidence that host × *Wolbachia* × environment interactions are pervasive — underscore the need for complementary field-based studies. Such work will deepen our understanding of *Wolbachia* spread and host effects in natural populations, while enabling more strategic selection of *Wolbachia* variants and host genotypes for deployment under specific applied contexts (Hoffmann and Cooper 2025).

## Materials & Methods

### Systems included in our study

We selected focal *Wolbachia* strains using the following criteria. First, strains were required to induce CI as reported in prior studies. Second, strains had to naturally inhabit *Drosophila* species that could be reared side-by-side on the same diet, enabling manipulation of temperature and cytotype (symbiotic vs aposymbiotic) while controlling for other environmental factors. Third, strains were strategically chosen to span a range of available genetic diversity, while including several closely related variants. This provided the opportunity for us to generalize our findings, while comparing closely and distantly related variants in different hosts. Based on these criteria, we selected eight *Drosophila* (*Wolbachia*) systems: *D. melanogaster* (*w*Mel), *D. teissieri* (*w*Tei), *D. seguyi* (*w*Seg), *D. chauvacae* (*w*Cha), *D. bicornuta* (*w*Bic), *D. triauraria* (*w*Tri), *D. simulans* (*w*Ri), and *D. simulans* (*w*Ha).

Our focal strains exhibit a range of CI phenotypes: *w*Tei causes weak-to-no CI, *w*Mel causes intermediate CI and male-age-dependent CI, and the remaining six strains cause relatively strong CI (Cooper et al. 2017; Shropshire et al. 2021a; Shropshire et al. 2022; Shropshire et al. 2026). Focal strains were collected from geographically diverse locations: *w*Tei from Bioko, *w*Seg from Cameroon, *w*Cha from Madagascar, *w*Bic from Taiwan, *w*Tri from Japan, *w*Ri from Riverside, California, and *w*Ha from Hawaii (O’Neill and Karr 1990; Turelli and Hoffmann 1991; Cooper et al. 2017; Turelli et al. 2018; Conner et al. 2021). The two *D. simulans* genotypes carry distinct mitochondrial haplotypes that co-transmit with their respective *Wolbachia*: *w*Ha is naturally associated with the siI haplotype and *w*Ri with siII (James and Ballard 2000; Ballard 2004).The *w*Mel strain is present in the *y*^*1*^*w*^*\**^ *Drosophila* line that was donated to the Bloomington *Drosophila* Stock Center (BDSC), a widely used reference strain for *Wolbachia* research due to its well-characterized CI phenotype (LePage et al. 2017; Shropshire et al. 2018; Layton et al. 2019; Shropshire and Bordenstein 2019; Shropshire et al. 2021a; Shropshire et al. 2022). While *D. melanogaster* is globally distributed (Richardson et al. 2012; Adrion et al. 2015), the original collection location of the *y*^*1*^*w*^*\**^ line is unknown. All lines were maintained in the laboratory since their collection at room temperature or in incubators at ~23 to 25°C.

Published genomic analyses reveal variation in *cif*-gene content and *Wovirus* composition among focal *Wolbachia*. The *cif* genes are classified into ten phylogenetic types (Types 1 to 10) based on their sequence divergence (LePage et al. 2017; Martinez et al. 2021; Amoros et al. 2025), while *Woviruses* are classified into four groups (sr1WO to sr4WO) based on serine recombinase alleles (Bordenstein and Bordenstein 2022). All eight focal strains encode at least one Type 1 *cif*-gene pair; *w*Ri carries two identical putatively disrupted copies (but see Shropshire et al. 2021b) and *w*Ha has a second copy that is putatively disrupted. Several strains carry additional *cif* types: Type 2 (*w*Bic and *w*Ri), Type 4 (*w*Tei), and Type 5 (*w*Tri) (Martinez et al. 2021; Shropshire et al. 2026). Regarding *Woviruses*, the sr3WO prophage is the most common, present in seven focal strains, either with one (*w*Tri, *w*Ha) or two (*w*Mel, *w*Tei, *w*Seg, *w*Cha, *w*Ri) copies. Additional *Wovirus* types include sr2WO (*w*Tei, *w*Tri, *w*Ri, and *w*Ha) and sr1WO (*w*Ri only) (Bordenstein and Bordenstein 2022; Shropshire et al. 2026). The *w*Bic genome has only a partial sr2WO-like serine recombinase sequence at the end of a contig, preventing further *Wovirus* characterization. Together, these focal *Wolbachia* differ in both their *cif*-gene repertoires and *Wovirus* complements, providing a basis for examining how these factors relate to CI strength.

### *Wolbachia* phylogenetics

We used the *Wolbachia* genomes listed in **Table S1** for phylogenetic reconstruction. Orthologous genes were identified using Prokka v.1.14.5 (Seemann 2014), and single-copy genes present in all genomes were extracted and aligned with MAFFT v.7 (Katoh and Standley 2013). To exclude pseudogenes and frameshifts, we excluded genes if they contained alignment gaps not in multiples of three or if gaps were present in more than one genome. For the phylogram presented in **Fig 1A** that included four *Nomada*-associated *Wolbachia* (*w*NLeu, *w*Nfa, *w*Npa, and *w*Nfe) as an outgroup to our eight focal strains, a total of 331 genes (285,540 bp) satisfied these criteria. To place *w*Cha of *D. chauvacae* among a larger set of *w*Mel-like *Wolbachia* presented in Fig S8 of Shropshire et al. (2026), we generated another phylogram with *w*Bic of *D. bicornuta* as an outgroup. For this set of 23 *Wolbachia*, a total of 333 genes (298,245 bp) satisfied our criteria, and we present the phylogram as a cladogram (**Fig S1**). We conducted phylogram inference under the GTR+Γ+I substitution model, partitioned by codon position. Each partition had an independent rate multiplier with prior Γ(1,1), with stationary frequencies and exchangeability rates drawn from flat, symmetrical Dirichlet distributions. We also inferred a relative chronogram for our set of eight *Wolbachia* plus the four *Nomada*-associated strains under the GTR+Γ model. We partitioned by codon position (with the above priors for each partition), fixed the relative root age to 1, and used the same birth-death prior as Turelli et al. (2018). A branch-rate prior of Γ(7,7) was normalized to a mean of 1 across all branches. We present the relative chronogram in **Fig. S8**. We completed four independent RevBayes runs for each tree and assessed the convergence for each run using Tracer (Rambaut et al. 2018). All four runs converged to the same result.

### Temperature-mapping and analyses

We obtained climatic data for *Wolbachia* collection sites from WorldClim v.2.1 (Fick and Hijmans 2017), which provides average monthly temperature for 1970–2000 at 30 arc-second resolution (~1 km at the equator). For each locale, three climatic variables were obtained: monthly minimum, average, and maximum temperatures. United States locations were analyzed at the state level (California and Hawaii) and Bioko island was isolated from mainland Equatorial Guinea using GADM level-1 administrative boundaries. All other locations were analyzed at the country level. We downloaded and processed these data using geodata v.0.0.6 (Hijmans 2025a) in R v.4.5.1 (R Core Team 2024). Country and state boundaries were obtained using rnaturalearth v.1.1.0 (Massicotte and South 2025). We processed and visualized spatial data using sf v.1.0.15 (Pebesma 2018), terra v.1.8.70 (Hijmans 2025b), and tidyterra v.0.7.2 (Hernangómez 2023). For each location, month, and variable, we calculated three metrics: mean across all pixels within the region, minimum value (coldest pixel), and maximum value (warmest pixel). We calculated annual average temperature as the mean across all 12 monthly averages. Spatial thermal heterogeneity was quantified as the difference between maximum and minimum pixel values for each temperature variable, and total temperature range was calculated as the difference between the warmest maximum and coldest minimum temperature across all months. Statistical comparisons were performed using Kruskal-Wallis tests followed by post-hoc Dunn’s tests with correction for pairwise comparisons. We created a global temperature map by averaging across all 12 monthly temperature layers and visualized temperature data using discrete color bins ranging from −55°C to 35°C (18 bins of 5°C each). Collection site coordinates were overlaid to show the geographic origin of focal strains. We refrain from directly corresponding site temperature estimates to observed trait variation since microclimatic variation and behavioral buffering — including at different life stages (Feder 1997; Gibbs et al. 2003) — influence the temperatures flies experience (Dillon et al. 2009).

### Insect lines, care, and maintenance

The following *Drosophila* (*Wolbachia*) lines were maintained: *D. bicornuta* (*w*Bic), *D. chauvacae* (*w*Cha), *D. melanogaster* (*w*Mel), *D. seguyi* (*w*Seg), *D. simulans* (*w*Ri), *D. simulans* (*w*Ha), *D. teissieri* (*w*Tei), and *D. triauraria* (*w*Tri). Aposymbiotic variants of each line were generated through tetracycline treatment following established protocols (protocol: Cooper and Shropshire 2024a). Treated lines were reared on standard food without antibiotics for a minimum of three generations to allow lines to recover prior to our experiments (Ballard and Melvin 2007). *Wolbachia* cytotype status was verified before and after experiments by PCR amplification of the *Wolbachia* surface protein (*wsp*) gene following a squish buffer DNA extraction protocol (protocol: Cooper and Shropshire 2024b). Stock information, sources, and PCR primers are provided in **Table S2** and **Table S3**.

All lines were reared on a standard cornmeal-based medium containing yellow cornmeal, dry corn syrup solids, malt extract, inactive dry yeast, soy flour, and agar, with tegosept and propionic acid added as mold and bacterial inhibitors (protocol: Wheeler et al. 2024). Stock populations were maintained at 23°C under a 12:12 h light:dark cycle with transfers to fresh vials every 2 to 3 weeks (protocol: Hartman et al. 2024). We supplemented all vials with a small strip of laboratory tissue (Kimberly-Clark KimWipes 34155) moistened with 0.5% propionic acid (Thermo Scientific 149300025) to prevent desiccation. For lines requiring higher humidity — particularly *D. teissieri* and *D. triauraria* at elevated temperatures — we placed a water tray on the bottom shelf of each incubator that was replenished as needed. Before generating experimental individuals, we maintained flies at 23°C for several generations. Once sufficient stock populations were established, flies were randomly distributed into four temperature conditions: 18°C, 20°C, 23°C, and 26°C. Approximately 20 to 25 unsexed, mated flies were added to vials containing 10 mL of food, and this process was repeated concurrently for each temperature. Flies laid eggs for 48 hours at the experimental temperature and were then transferred to fresh vials. This process was repeated until sufficient vials were available for virgin collection. Virgins were collected under CO_2_ anesthesia.

### Hatch-rate analyses

All crosses used 3- to 4-day-old virgin females and 0- to 1-day-old virgin males. We reared males from egg to adult at experimental temperatures (18°C, 20°C, 23°C, or 26°C), while females were maintained at 23°C until crossing. Although we collected males with and without *w*Tei at all four temperatures, females did not lay eggs at 18°C. Males with and without *w*Tri were collected at only two temperatures (23°C and 26°C) due to the two coolest temperatures inhibiting *D. triauraria* development. Three cross types assessed CI: aposymbiotic males × aposymbiotic females (compatible control), symbiotic males × aposymbiotic females (CI), and symbiotic males × symbiotic females (rescue). Mating pairs were placed in vials containing an ice-cream spoon filled with fly food that we supplemented with blue food coloring, 0.1 g of extra agar per 100 mL of food, and fresh yeast (protocol: Shropshire 2025). Vials were incubated overnight at the experimental temperature. We then transferred flies to new vials with fresh spoons, a process that we repeated for five consecutive days. Immediately after fly removal, we counted all embryos on each spoon (total eggs). After 48 hours of incubation at the experimental temperature, we counted unhatched embryos. Hatched embryos were calculated as total eggs minus unhatched embryos. Hatch rate was calculated as the proportion of embryos hatched per replicate. Replicates with fewer than 10 embryos were excluded to ensure statistical reliability (Shropshire et al. 2022). During egg-lay experiments, flies were transferred between vials using mouth aspiration to avoid CO_2_ exposure (Shropshire et al. 2021a; Shropshire et al. 2022).

We analyzed embryo hatch rates using a zero-inflated binomial generalized linear mixed model (GLMM) with a logit link function. The response variable was specified as a two-column matrix (hatched, total − hatched), and the model included strain (eight levels), temperature (18°C, 20°C, 23°C, 26°C), and cross type (compatible, CI, rescue) as fixed effects with all two- and three-way interactions. An observation-level random effect accounted for overdispersion, and the zero-inflation component was modeled with an intercept-only term. Models were fitted using glmmTMB v.1.1.13 (McGillycuddy et al. 2025). We assessed significance of fixed effects using Type II ANOVA with car v.3.1.3 (Fox and Weisberg 2019) and evaluated model fit using DHARMa v.0.4.7 (Hartig 2024), including tests for residual normality (Kolmogorov-Smirnov test), dispersion, outliers, and zero-inflation.

We calculated estimated marginal means for each strain × temperature × cross type combination using emmeans v.1.11.2.8 with type = “response” to obtain predicted probabilities (Lenth 2025). CI strength was quantified as the odds ratio (*OR*_CI_) comparing hatching odds in CI crosses to compatible crosses. Rescue efficiency was quantified as the odds ratio (*OR*_R_) comparing rescue crosses to compatible crosses. Confidence intervals and *P*-values used FDR correction for multiple comparisons. We evaluated temperature effects on CI strength within each strain by comparing *OR*_CI_ values across temperatures; the ratio of odds ratios (*OR*_CI,T_) was calculated for each pairwise temperature comparison. To quantify the magnitude of temperature effects within each strain, we fitted per-strain binomial GLMMs (compatible and CI crosses only); zero-inflated models were attempted first, with non-zero-inflated fallback if convergence failed (*w*Mel, *w*Cha, *w*Tri). Partial η^2^ was calculated from the cross × temperature interaction: partial η^2^ = χ^2^ / (χ^2^ + n).

### Developmental timing

We measured development time for a subset of *Wolbachia*-bearing and aposymbiotic lines. Approximately 50 mated adults (4- to 5-day-old) were placed in 6 oz *Drosophila* bottles fitted with inverted 35 mm Petri dishes containing 15 mL of standard fly food, secured with tape. Assemblies were placed in incubators at experimental temperatures (18°C, 20°C, 23°C, or 26°C). Adult females oviposited directly onto the food surface over a 48-hour egg-lay window. After this window, a thin layer of food containing eggs was carefully sliced from each plate. Eggs were visualized and counted under a Leica S9i stereomicroscope. Approximately 80 to 100 eggs were transferred to individual food vials containing fresh fly food. Development vials were not supplemented with KimWipes containing 0.5% propionic acid; instead, we gently sprayed 0.5% propionic acid onto the food surface every 3 to 4 days. Vials were monitored every other day beginning on the day of egg transfer. The date of first adult emergence in each vial was recorded. We defined development time as the number of days between egg transfer and first adult emergence.

We analyzed development time data using permutation-based approaches to account for zero within-group variance in several strain-temperature-cytotype combinations. An omnibus test was performed using permutation-based multivariate analysis of variance (PERMANOVA) with vegan v.2.7.2 (Oksanen et al. 2025). The model included system (six *Wolbachia*-host systems), cytotype (aposymbiotic, symbiotic), temperature (18°C, 20°C, 23°C, 26°C), and all two- and three-way interactions. PERMANOVA used 9,999 permutations, Euclidean distance, and Type I (sequential) sums of squares. Pairwise permutation tests (10,000 permutations each) evaluated cytotype effects within each strain-temperature combination and temperature effects within each strain-cytotype combination. We calculated *P*-values as the proportion of permuted differences with absolute values ≥ the observed difference. When no permutation exceeded the observed value, *P* = 1/(n + 1) ≈ 9.9 × 10^−5^. All *P*-values were FDR-corrected, with significance assessed at α = 0.05. For groups with non-zero variance, we estimated 95% confidence intervals by bootstrap resampling (10,000 samples) using boot v.1.3.32.

### Tissue collection

We dissected testes from male siblings of flies used in hatch rate assays to quantify *Wolbachia* densities, *Wovirus* densities, and *cif*-gene transcript levels. Tissue collections were synchronized with crossing experiments: dissections occurred the day after the majority of egg-lay replicates exceeded 10 embryos for each strain × temperature combination. This timing ensured that molecular measurements reflected the physiological state of males when CI-affected eggs were laid. We performed dissections under a stereomicroscope (Leica S9i) in ice-cold RNase-free phosphate-buffered saline (MilliporeSigma OmniPur 6507-4L). For each biological replicate, we removed both testes from a single male, cleaned them of adhering tissue, and transferred them to a 1.5 mL microcentrifuge tube. To minimize RNA degradation, dissection tools and working surfaces were cleaned with RNase AWAY (Thermo Scientific 7002) between samples. Samples for DNA extraction were collected from all eight strains. Each replicate comprised five pooled pairs of testes in tubes containing three 3 mm borosilicate glass beads. We collected samples for RNA extraction from *w*Mel, *w*Ri, and *w*Ha-carrying males only, with each replicate comprising five pooled pairs of testes in tubes containing 400 μL TRIzol Reagent (Invitrogen 15596018) and three 3 mm glass beads. All samples were homogenized using a TissueLyser II bead mill (Qiagen 85300) at 25 Hz for 2 min, pulse-centrifuged, and stored at −80°C.

### *Wolbachia* and *Wovirus* densities

DNA samples were thawed and further homogenized using a TissueLyser II at 25 Hz for 2 min. We extracted DNA using the DNeasy Blood and Tissue Kit (Qiagen 69506) (protocol: Hartman and Shropshire 2024), storing purified DNA at 4°C for short-term storage or −80°C for long-term preservation. *Wolbachia* and *Wovirus* densities were quantified by qPCR. We conducted all reactions in triplicate in 10 μL volumes using PowerUp SYBR Green Master Mix (Applied Biosystems A25742) at 1× concentration. Three target genes were amplified: the *Drosophila* gene nAcRalpha-34E (Hague et al. 2024), the *Wolbachia* gene *ftsZ* (Shropshire et al. 2021a; Shropshire et al. 2022), and the *Wovirus* serine recombinase gene (Bordenstein and Bordenstein 2022). A single primer pair amplified nAcRalpha-34E and *ftsZ* across systems, while multiple primers were required for the *Wovirus* serine recombinase (**Table S3**). Cycling conditions were 50°C for 2 min, 95°C for 2 min, followed by 40 cycles of 95°C for 15 s and 60°C for 1 min. We calculated *Wolbachia* density using the ΔC_q_ method: ΔC_q_(*Wolbachia*) = C_q_(*ftsZ*) – C_q_(nAcRalpha-34E), with fold-change calculated as 2^−ΔCq^. This provides a measure of *Wolbachia* gene copies relative to host gene copies. *Wovirus* density was calculated similarly but relative to *Wolbachia*: ΔC_q_(*Wovirus*) = C_q_(serine recombinase) – C_q_(*ftsZ*), with fold-change = 2^−ΔCq^. For strains with multiple *Wovirus* variants detected by different primer sets (**Table S3**), total *Wovirus* density was calculated by summing fold-changes across all detected variants.

Log_2_-transformed *Wolbachia* and *Wovirus* densities were analyzed using linear models with *Wolbachia* strain (eight focal strains), temperature (18°C, 20°C, 23°C, and 26°C), and their interaction as fixed effects. We assessed significance of main effects and interactions using Type II ANOVA with car v.3.1.3 (Fox and Weisberg 2019). Model validation was performed using DHARMa v.0.4.7 diagnostics (Hartig 2024) following conversion to glmmTMB v.1.1.13 format (McGillycuddy et al. 2025), testing for residual normality (Kolmogorov-Smirnov test), dispersion, outliers, and zero-inflation. Partial ω^2^ values quantified effect sizes for each model term. We conducted pairwise comparisons of estimated marginal means across temperatures within each strain using emmeans v.1.11.2.8 (Lenth 2025) with FDR correction. To quantify the magnitude of temperature effects within each strain, we fitted per-strain linear models and calculated partial ω^2^ from the temperature term. Furthermore, to evaluate the relationship between *Wolbachia* and *Wovirus* densities within each strain, we calculated Person correlation coefficients between log_2_-transformed *Wolbachia* densities and log_2_-transformed *Wovirus* densities. Correlations were computed separately for each of the six strains with detectable *Wovirus*.

### Gene expression assays

RNA samples were thawed, supplemented with an additional 400 μL of TRIzol Reagent, and further homogenized using a TissueLyser II at 25 Hz for 2 min. To control for variation in RNA extraction efficiency, we added 1 μL of TATAA Universal RNA Spike I (TATAA Biocenter RS25SI) at 1:16 dilution to each sample after homogenization. RNA was extracted and purified via TRIzol:chloroform phase separation (protocol: Van Vlaenderen and Shropshire 2025; Van Vlaenderen et al. 2026). To remove residual genomic DNA, we treated purified RNA with DNase using the Invitrogen DNA-free kit following the manufacturer’s “rigorous” treatment protocol. Successful DNA removal was confirmed by PCR amplification of the host 28S rRNA gene, which yielded no product. We performed first-strand cDNA synthesis using SuperScript IV VILO Master Mix following the manufacturer’s protocol. cDNA was stored at 4°C for short-term use or −80°C for long-term preservation.

We quantified *cifB*-transcript abundance using reverse transcription-digital droplet PCR (RT-ddPCR) on cDNA derived from testes of males carrying *w*Mel, *w*Ri, and *w*Ha. We used three primer-probe sets with FAM-labeled probes to capture all *cifB* variants across the three focal strains (**Table S3**), one of which was characterized in detail with a limit of detection of 3 *cifB* copies per reaction (Van Vlaenderen et al. 2026). We calculated transcript abundance as the concentration of *cifB* transcripts (copies/μL) divided by the concentration of spike-in transcripts (copies/μL), providing a measure normalized for RNA purification and processing efficiency.

We analyzed log_2_-transformed *cifB*-transcript abundance using a linear model with *cifB* variant (five variants), temperature treatment (cool vs warm), and their interaction as fixed effects. Cool temperatures were 18°C for *w*Mel and *w*Ri and 20°C for *w*Ha; warm was 23°C for all strains. We assessed significance using Type II ANOVA with car v.3.1.3 (Fox and Weisberg 2019). Model validation used DHARMa v.0.4.7 diagnostics (Hartig 2024) following conversion to glmmTMB v.1.1.13 format (McGillycuddy et al. 2025). Partial ω^2^ values quantified effect sizes. Pairwise comparisons between cool and warm temperatures within each variant used emmeans v.1.11.2.8 (Lenth 2025) with FDR correction. We calculated fold-change in *cifB*-transcript abundance between cool and warm temperatures from estimated marginal means. Fold-change was calculated as 2^Δ^, where Δ is the difference in estimated marginal means on the log_2_ scale.

### Within-strain concordance analysis

We evaluated within-strain concordance between temperature-induced changes in different variables using Kendall’s tau (τ). Uncertainty in estimated marginal means was propagated to τ estimates using Monte Carlo simulations (*N* = 10,000 iterations). For each iteration, we sampled values from Normal(estimated marginal means, SE^2^) distributions for each condition and computed τ from the sampled values. We calculated point estimates by averaging Monte Carlo samples on the Fisher z-transformed scale, then back-transforming to the correlation scale. Probability of direction (*p*_d_) was calculated as the proportion of Monte Carlo samples with the same sign as the point estimate; this metric quantifies directional certainty. We evaluated concordance for the following within-strain relationships: CI strength vs development time, CI strength vs *Wolbachia* density, CI strength vs *cifB*-transcript abundance, *Wolbachia* density vs *Wovirus* density, and *cifB*-transcript abundance vs *Wolbachia* density.

### Bayesian phylogenetic regression models

Bayesian phylogenetic mixed models were used to evaluate relationships between variables while accounting for phylogenetic non-independence among *Wolbachia* strains. All models were fitted using brms v.2.23.0 (Bürkner 2017) with a phylogenetic covariance matrix derived from our relative *Wolbachia* chronogram (**Fig S8**). Models were fitted with Gaussian likelihood using Hamiltonian Monte Carlo sampling with 4 chains of 8,000 iterations each (2,000 warmup; 24,000 post-warup draws total) and adapt_delta = 0.99 to minimize divergent transitions. We assessed convergence by verifying that all R^A^ values were < 1.01. For each model, results reported include the posterior mean of the slope (β), 95% credible interval, Bayesian R^2^, and probability of direction (*p*_d_). The probability of direction indicates the proportion of the posterior distribution consistent with the sign of the point estimate.

Three sets of relationships were evaluated using the general formula: response ~ predictor + (1 | gr(Strain, cov = A)), where A is the phylogenetic covariance matrix. First, we modeled CI strength (log *OR*_CI_) as a function of *Wolbachia* density, development time, or *cifB*-transcript abundance using estimated marginal means; these models were fitted for all strains, temperature-sensitive strains (*w*Mel, *w*Tei, *w*Ri, *w*Ha), and temperature-resistant strains (*w*Seg, *w*Cha, *w*Bic, *w*Tri) separately, with the exception of CI strength ~ *cifB*, which we modeled only for *w*Mel, *w*Ri, and *w*Ha. Second, we modeled *cifB* transcription as a function of *Wolbachia* density using estimated marginal means from strains with matched measurements (*w*Mel, *w*Ri, *w*Ha). Third, we modeled *Wolbachia* density as a function of *Wovirus* density using raw log_2_-transformed measurements from the six strains with *Wovirus* data; this model included an additional temperature random effect.

## Acknowledgments

We thank Kelley Van Vaerenbergh and Erin Markham for assistance with assay optimization and collection of preliminary data not included in this study. We are grateful to Lara Parada-Tixe and Abby Klebe for their assistance with laboratory maintenance in the Shropshire Lab. Finally, we thank all members of the Cooper and Shropshire Labs for their discussions.

## Figure generation

We generated all figures in R version 4.5.1 (R Core Team 2024) using ggplot2 (Wickham 2016). Final figure aesthetics were refined using Inkscape v1.4.2.

## Data availability

All data are openly available in the Dryad Digital Repository at https://doi.org/10.5061/dryad.18931zdbf (peer review link: [here](http://datadryad.org/share/LINK_NOT_FOR_PUBLICATION/0VmvhcVpcdiYC9BsL3Z8110mI8jQYIU1L_WlrDhySHU)). Scripts used for data analysis are available on GitHub at https://github.com/JDShropshire/Temperature-Sensitive-CI.

