## Supplementary Material for "Temperature-sensitive cytoplasmic incompatibility across divergent *Wolbachia* partly reflects *cifB* transcription, not endosymbiont density"

#### Supporting Results

##### Focal *Wolbachia-Drosophila* genotypes were sampled from thermally diverse locations

Of the eight focal systems, the country (or US state) of origin is known. Of these seven locations, Cameroon (*w*Seg; 25.5°C annual average) is the warmest, followed by Madagascar (*w*Cha; 22.9°C), Bioko island of Equatorial Guinea (*w*Tei; 22.4°C), Hawaii (*w*Ha; 18.8°C), Taiwan (*w*Bic; 18.8°C), California (*w*Ri; 14.0°C), and Japan (*w*Tri; 7.0°C) (**Fig 1B**; **Fig S2A**). Monthly average temperature in Japan is cooler than all other locations ($P < 0.05$); California is cooler than Cameroon, Madagascar, and Bioko ($P < 0.05$); and Cameroon, Madagascar, Bioko, Taiwan, and Hawaii are statistically similar ($P > 0.05$). However, average temperatures vary spatially across sites within each country and state. The range between the coolest and warmest average temperatures within each location is highest in Japan (−24.4°C to 29.7°C, 54.1°C range) and progressively becomes more stable in California (−10.7°C to 37.9°C, 48.6°C range), Taiwan (−1.4°C to 29.8°C, 31.2°C range), Cameroon (6.1°C to 34.0°C, 27.9°C range), Hawaii (1.3°C to 27.2°C, 25.9°C range), Madagascar (7.4°C to 30.0°C, 22.6°C range), and Bioko (10.6°C to 26.5°C, 15.9°C range). Average temperature is spatially more stable in Bioko and Madagascar and less stable in Japan than all other locations ($P < 0.05$; **Fig S2B**). Variations in average high and low temperatures, and their spatial distribution across each location, follow similar trends, as does the range from the lowest low to highest high in each location (**Fig S2C–G**). The exact collection sites within these regions are generally unknown, preventing finer-scale temperature analysis. We refrain from directly corresponding site temperature estimates to observed trait variation since microclimatic variation and behavioral buffering, including at different life stages (Feder 1997; Gibbs et al. 2003), influence the temperatures flies experience (Dillon et al. 2009).

##### Temperature affects embryo hatching from compatible crosses in four focal systems

Before analyzing temperature effects on CI, we first assessed how temperature influences hatch rates in compatible crosses. We calculated the estimated marginal means from the GLMM, which provides model-predicted egg-hatch rates for each system at each temperature. Temperature did not significantly affect compatible crosses in *D. chauvacae* ($\text{min}_{26^\circ\text{C}}$ = 0.95 [0.80 to 0.99], $\text{max}_{23^\circ\text{C}}$ = 0.99 [0.95 to 1.0], $P = 0.31$), *D. triauraria* ($\text{min}_{23^\circ\text{C}}$ = 0.92 [0.75 to 0.98], $\text{max}_{26^\circ\text{C}}$ = 0.97 [0.88 to 0.99], $P = 0.27$), or in the *D. simulans* genotype hosting *w*Ha ($\text{min}_{20^\circ\text{C}}$ = 0.94 [0.89 to 0.97], $\text{max}_{23^\circ\text{C}}$ = 0.99 [0.95 to 1.0], $P = 0.32$; **Fig S3**). In *D. melanogaster* ($\text{min}_{18^\circ\text{C}}$ = 0.86 [0.68 to 0.95], $\text{max}_{20^\circ\text{C}}$ = 0.98 [0.94 to 0.99], $P = 0.08$) and *D. seguyi* ($\text{min}_{18^\circ\text{C}}$ = 0.77 [0.51 to 0.92], $\text{max}_{26^\circ\text{C}}$ = 0.97 [0.90 to 0.99], $P = 0.058$) differences approached, but did not reach, statistical significance ($\alpha = 0.05$; **Fig S3**). Notably, we were unable to successfully rear *D. triauraria* at 18 or 20°C, highlighting significant negative effects of the coolest temperatures on this species, which precluded analysis of downstream egg hatch. In contrast, the egg hatch of compatible crosses in *D. teissieri* varied significantly across temperatures, with the lowest hatch observed at 26°C (0.085 [0.018 to 0.32]) and the highest at 23°C (0.98 [0.90 to 1.0], $P = 1.1e\text{-}6$). Compatible crosses in *D. bicornuta* ($min_{26°C}$ = 0.89 [0.74 to 0.96], $max_{23°C}$ = 0.99 [0.98 to 1.0], $P = 4.9e\text{-}3$) and in the *D. simulans* genotype hosting *w*Ri ($min_{18°C}$ = 0.94 [0.85 to 0.98], $max_{26°C}$ = 1.0 [0.99 to 1.0], $P = 1.6e\text{-}3$; **Fig S3**) showed relatively small, but significant, variation in egg hatch across temperatures. In summary, the egg hatch of compatible crosses is temperature-sensitive in some systems, and temperature-resistant in others.

**Temperature strongly modulates development time across all systems**

Given temperature's dominant influence on development time, we characterized temperature effects within each strain-cytotype combination using pairwise permutation tests across all temperature pairs. All testable strain-cytotype combinations exhibit significant temperature-sensitive development time variation ($P < 0.05$ for at least one comparison within each group). Development time decreases monotonically with increasing temperature in all systems. For symbiotic vials, *D. melanogaster* development time decreases from 18.0 days at 18°C to 13.1 days at 20°C, 9.6 days at 23°C, and 9.0 days at 26°C, with all pairwise comparisons significant ($P < 0.05$; **Fig S5**). Similar patterns are observed across symbiotic *D. teissieri* (9.3 days at 20°C to 8.0 days at 26°C), *D. chauvacae* (18.0 days at 18°C to 12.0 days at 26°C), *D. bicornuta* (19.6 days at 18°C to 8.0 days at 26°C), and *D. simulans* that harbors *w*Ri (15.4 days at 18°C to 8.0 days at 26°C; **Fig S5**). The magnitude of temperature effects varies among strains, with *w*Bic exhibiting the largest absolute change (11.6 days between 18°C and 26°C). Notably, several groups exhibit zero variance at specific temperatures, where all observations are identical (*e.g., D. melanogaster* at 26°C, *D. teissieri* at 23°C and 26°C), reflecting extreme developmental synchrony under those conditions. Aposymbiotic flies show similar temperature-sensitive trends (**Fig S5**). These results demonstrate that temperature exerts profound effects on development time across focal systems.

### Supporting Information

#### Table S1. *Wolbachia* genomes used in this study.

| **Host (*Wolbachia*)** | ***Wolbachia* accession** |
| --- | --- |
| *Drosophila melanogaster* (*w*Mel) | GCF_000008025.1 |
| *Drosophila teissieri* (*w*Tei) | GCA_018689955.1 |
| *Drosophila seguyi* (*w*Seg) | GCA_032584075.1 |
| *Drosophila chauvacae* (*w*Cha) | JAPQLI000000000.1 |
| *Drosophila bicornuta* (*w*Bic) | GCF_028982105.1 |
| *Drosophila triauraria* (*w*Tri) | GCA_014129515.1 |
| *Drosophila simulans* (*w*Ri) | GCF_000022285.1 |
| *Drosophila simulans* (*w*Ha) | CP003884.1 |
| *Nomada leucophthalma* (*w*NLeu) | GCA_001675715.1 |
| *Nomada flava* (*w*NFla) | GCA_001675695.1 |
| *Nomada panzeri* (*w*NPa) | GCA_001675775.1 |
| *Nomada ferruginata* (*w*NFe) | GCA_001675785.1 |
| *Drosophila bocqueti* (*w*Bocq) | GCF_032584155.1 |
| *Drosophila sp. aff. chauvacae* (*w*Ach) | GCF_032848195.1 |
| *Drosophila tristis* (*w*Tris) | GCF_028981745.1 |
| *Drosophila recens* (*w*Rec) | GCA_000742435.1 |
| *Drosophila seguyi* (*w*Seg) | GCA_032584075.1 |
| *Drosophila malagassya* (*w*Mal) | GCA_032584035.1 |
| *Drosophila santomea* (*w*San) | GCA_018467135.1 |
| *Drosophila yakuba* (*w*Yak) | GCA_018467115.1 |
| *Zaprionus taronus* (*w*Zta) | GCF_033042245.1 |
| *Zaprionus tsacasi* (*w*Zts) | GCF_032849005.1 |
| *Drosophila borealis* (*w*Bor) | GCA_014129615.1 |
| *Drosophila innubila* (*w*Inn) | GCA_021378375.1 |
| *Drosophila simulans* (*w*Au) | GCA_017916175.1 |
| *Drosophila tropicalis* (*w*Tro) | GCA_014129525.1 |
| *Drosophila incompta* (*w*Inc) | GCA_001758565.1 |
| *Drosophila arawakana* (*w*Ara) | GCA_014129655.1 |
| *Scaptomyza pallida* (*w*Spa) | GCA_032584115.1 |
| *Sphyracephala brevicornis* (*w*Sbr) | GCA_902647005.1 |
| *Diachasma alloeum* (*w*Dal) | GCA_902646845.1 |

**Table S2. *Drosophila* lines used in this study.** “Tet” in the Line column indicates that the line following the colon was treated with tetracycline to generate an aposymbiotic line.

| Host | Cytotype | Line |
|---|---|---|
| *D. melanogaster* | *w*Mel | wMel_yw |
| *D. melanogaster* | aposymbiotic | Tet: wMel_yw |
| *D. teissieri* | *w*Tei | wTei_B13_L11 |
| *D. teissieri* | aposymbiotic | Tet: wTei_B13_L11 |
| *D. seguyi* | *w*Seg | wSeg_L1 |
| *D. seguyi* | aposymbiotic | Tet: wSeg_L1 |
| *D. chauvacae* | *w*Cha | wCha_chavaucea_L1 |
| *D. chauvacae* | aposymbiotic | Tet: wCha_chavaucea_L1 |
| *D. bicornuta* | *w*Bic | wBic |
| *D. bicornuta* | aposymbiotic | Tet: wBic |
| *D. triaruaria* | *w*Tri | wTri_L1 |
| *D. triauraria* | aposymbiotic | Tet: wTri_L1 |
| *D. simulans* | *w*Ri | wRi_Riv_84 |
| *D. simulans* | aposymbiotic | Tet: wRi_Riv_84 |
| *D. simulans* | *w*Ha | wHa_SIM_CAR5 |
| *D. simulans* | aposymbiotic | Tet: wHa_SIM_CAR5 |

#### Table S3. Primers used in this study.

| Target | Forward sequence (5'-3') | Reverse sequence (5'-3') | Probe sequence (5'-3') | Purpose |
|---|---|---|---|---|
| *wsp* | TGGTCCAATAAGTGATGAAGAAAC | AAAAATTAAACGCTACTCCA | | PCR for *Wolbachia* (all strains) |
| *28s* | TACCGTGAGGGAAAGTTGAAA | AGACTCCTTGGTCCGTGTTT | | PCR for host (all species) |
| *ftsZ* | GCAGCCAATAGAGTGCGTG | TTCCCTCCATCGCTTGATCA | | qPCR for *Wolbachia* (all strains) |
| *nAcRalpha-34E* | CTATGGTCGTTGACAGACT | GTAGTACAGCTATTGTGGC | | qPCR for host (all species) |
| sr1WO (*w*Ri) | AAACTTCATGCGGCCAAAGC | TCTTTCTTGGCCAACCCGAC | | qPCR for serine recombinase |
| sr2WO (*w*Ri, *w*Tri) | GCCAAATTGCCGAGCTCAAG | CGCTTCTAAACCTTCACGCC | | qPCR for serine recombinase |
| sr2WO (*w*Tei) | ATGACCAGGTAGAAGTGGCG | TGACGTCCAGTACAATGTTGC | | qPCR for serine recombinase |
| sr3WO (*w*Mel 2, *w*Tei 2, *w*Seg 2, *w*Ha), sr2WO (*w*Ha) | AGTCTTGATGCACAGCGAGT | TTGGCTAATGCTACCCACCC | | qPCR for serine recombinase |
| sr3WO (*w*Mel 1, *w*Ri 1/2, *w*Tri) | TGCTGAAAGAGGTAAGATGCG | AATGCTACCCACCCTTCTCG | | qPCR for serine recombinase |
| sr3WO (*w*Tei 1, *w*Seg 1) | ATATGCTTCACTGTCCTTGCTTC | CAAACGGTATTGTCAGGAGCAG | | qPCR for serine recombinase |
| sr3WO (*w*Cha) | CGCAACGAGTAGCATGTGAG | ATAGCCGCCATCATCGTACC | | qPCR for serine recombinase |
| *cifBw*Mel[T1], *cifBw*Ri[T1], *cifBw*Ha[T1-1] | GCAAGGTACTAGAGCACAGG | CACGAGCGTTGTTTCTACG | AGGTGGTACTTCTACAGCACAAGG | RT-ddPCR for *cifB* |
| *cifBw*Ha[T1-2] | GGTGGATGGAGATCTTGAAGG | AATACAGGGAGGAAACGACC | TGTCCATCTTTCTTGTATCATGTAAGTGCA | RT-ddPCR for *cifB* |
| *cifBw*Ri[T2] | CGTTAGTAATGTGCATCGCG | AACCGTTCATGACTGTCTGG | TGCAAATGTAATGATGAATACTGGCTGGG | RT-ddPCR for *cifB* |

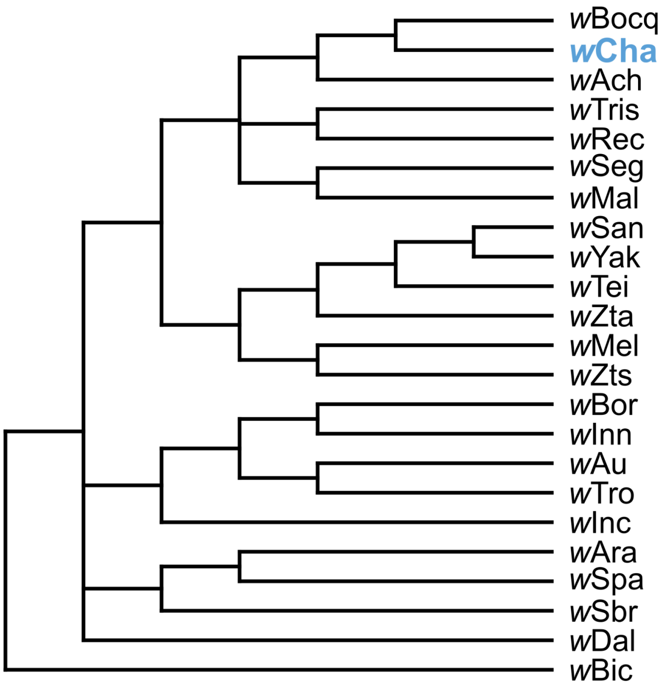


**Figure S1. Phylogenetic placement of *w*Cha among *w*Mel-like *Wolbachia*.** A cladogram based on 333 genes (298,245 bp) with a bifurcating history (see Shropshire et al. 2026). *Wolbachia* include *w*Cha of *D. chavacae*, the *w*Mel-like *Wolbachia* from Fig S8 in Shropshire et al. (2026), and outgroup *w*Bic of *D. bicornuta*. These *w*Mel-like *Wolbachia* diverged about 889 KYA and are observed in dipteran and hymenopteran hosts that diverged approximately 350 MYA, and that span the Holometabola (Wang et al. 2016). Nodes with posterior probability < 0.95 were collapsed into polytomies. The analysis places *w*Cha as a member of the *w*Mel-like *Wolbachia* clade that is most closely related to *w*Bocq from *D. bocqueti*.

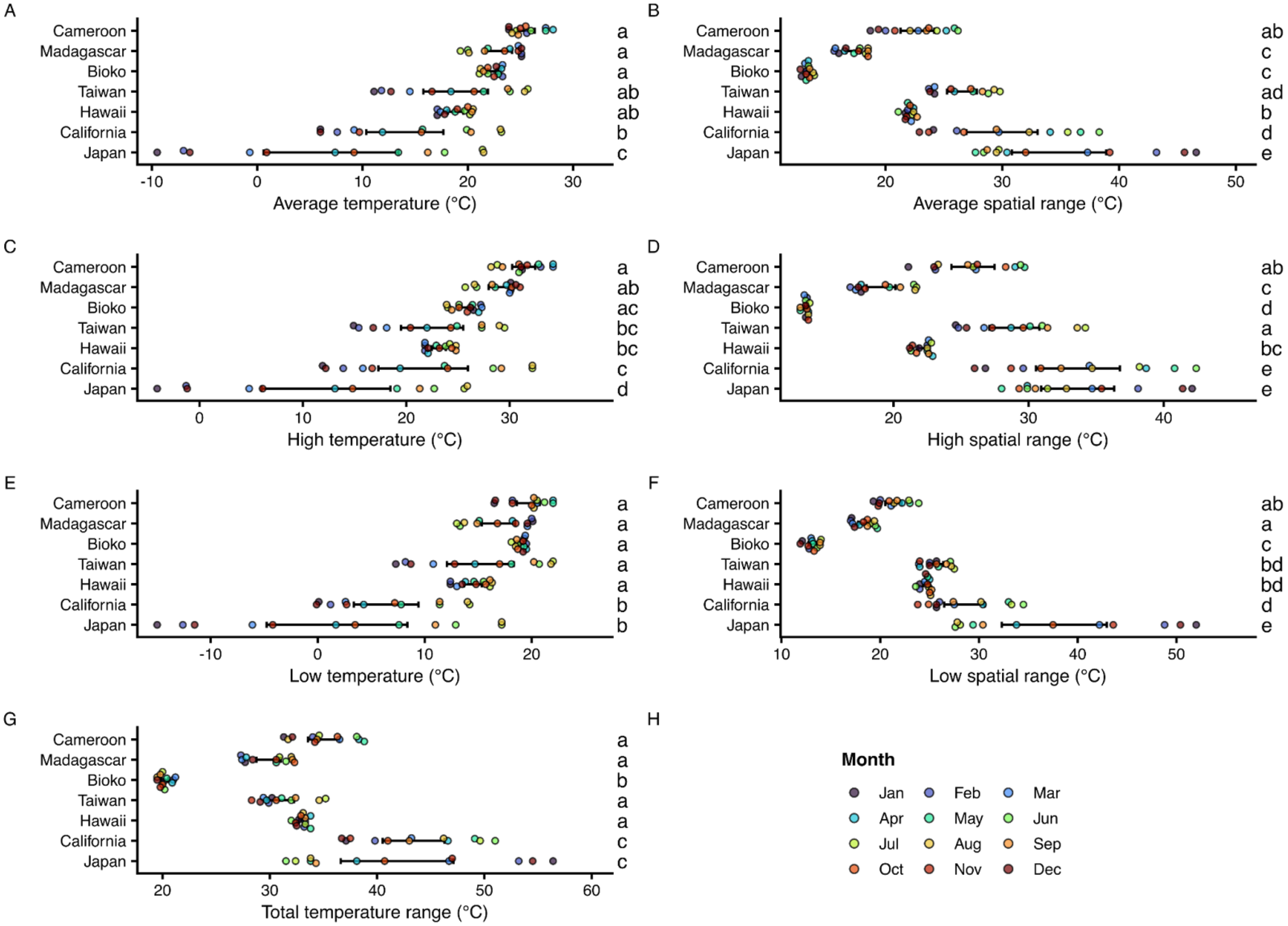


**Figure S2. Temperature distributions differ significantly among collection regions. (A)** Average temperature represents the mean temperature across all pixels within each region. **(B)** Average spatial range quantifies temperature variation within each region, calculated as the difference between the warmest and coldest pixels. **(C)** High temperature represents the monthly maximum temperature averaged across each region. **(D)** High spatial range shows variation in maximum temperatures within regions. **(E)** Low temperature represents the monthly minimum temperature averaged across each region. **(F)** Low spatial range shows variation in minimum temperatures within regions. **(G)** Total temperature range represents the difference between the absolute warmest maximum and coldest minimum temperatures within each region across all months. Locations are ordered by average annual temperature from warmest (Cameroon) to coldest (Japan). Colored points represent individual months ($N$ = 12 per location). Error bars show 95% confidence intervals. Spatial range metrics quantify temperature variation across pixels within each region. Letters indicate significant differences in CI strength observed among temperatures within each strain after Tukey's Honest Significant Difference post-hoc tests ($\alpha$ = 0.05); shared letters denote non-significant differences. Temperature data are from WorldClim 2.1 at 30 arc-second resolution (~1 km at the equator, 1970-2000 averages).

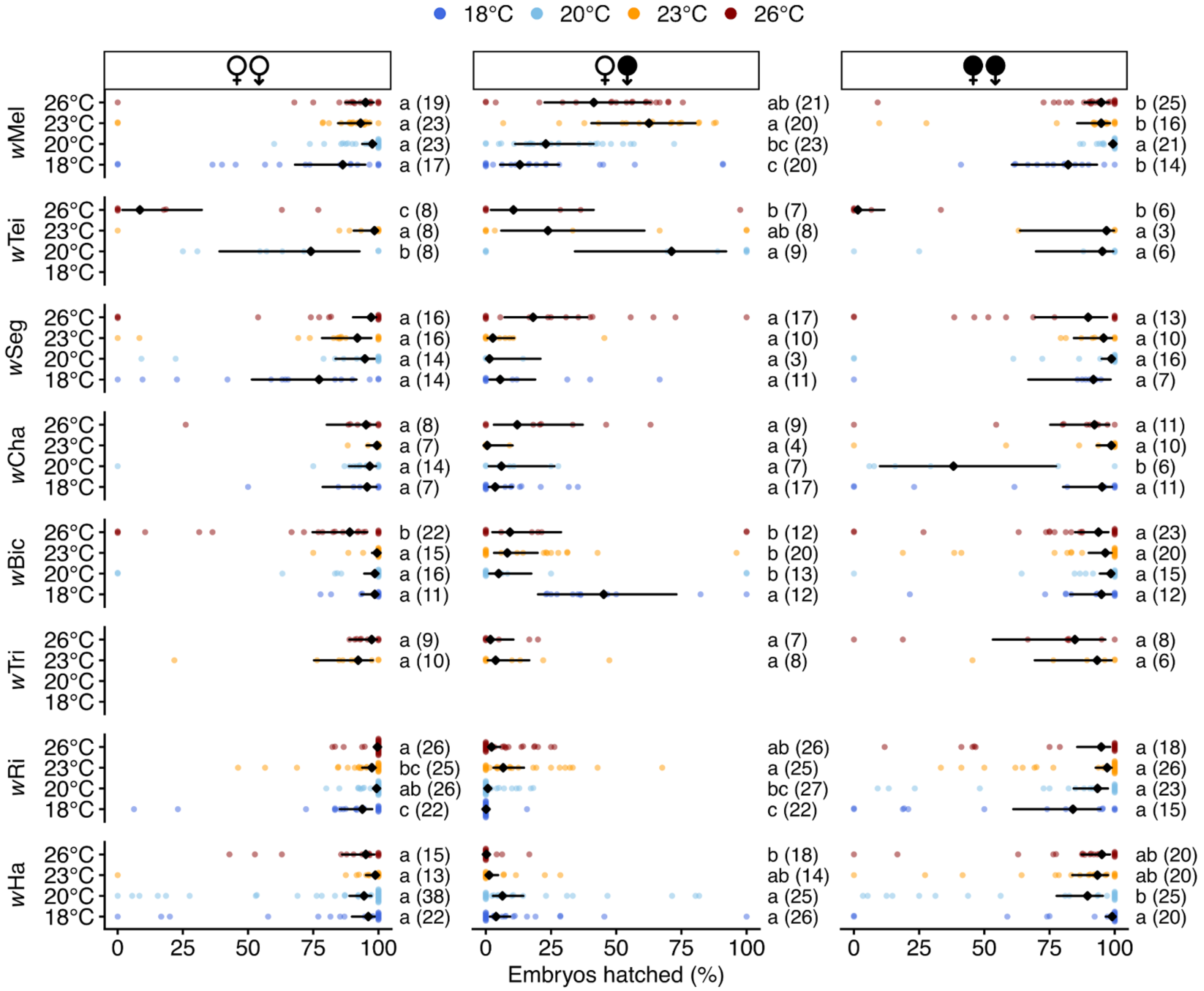


**Figure S3. Temperature affects embryonic hatching in strain- and cross-specific patterns.** Embryonic hatch rates from mating pairs. Column facets represent cross types indicated by female (♀) and male (♂) symbols: filled symbols indicate *Wolbachia*-bearing (symbiotic) individuals while unfilled symbols indicate *Wolbachia*-free (aposymbiotic) individuals. Row facets represent *Wolbachia* strain-host combinations. Y-axis shows experimental temperatures (18°C, 20°C, 23°C, 26°C). Individual data points (colored dots) show the percentage of embryos that hatched from each cross. Black diamonds represent estimated means with error bars depicting 95% confidence intervals from a zero-inflated binomial generalized linear mixed model. Letters indicate significant differences in CI strength observed among temperatures within each strain (FDR-corrected, $\alpha = 0.05$); shared letters denote non-significant differences. Sample sizes per group are in parentheses.

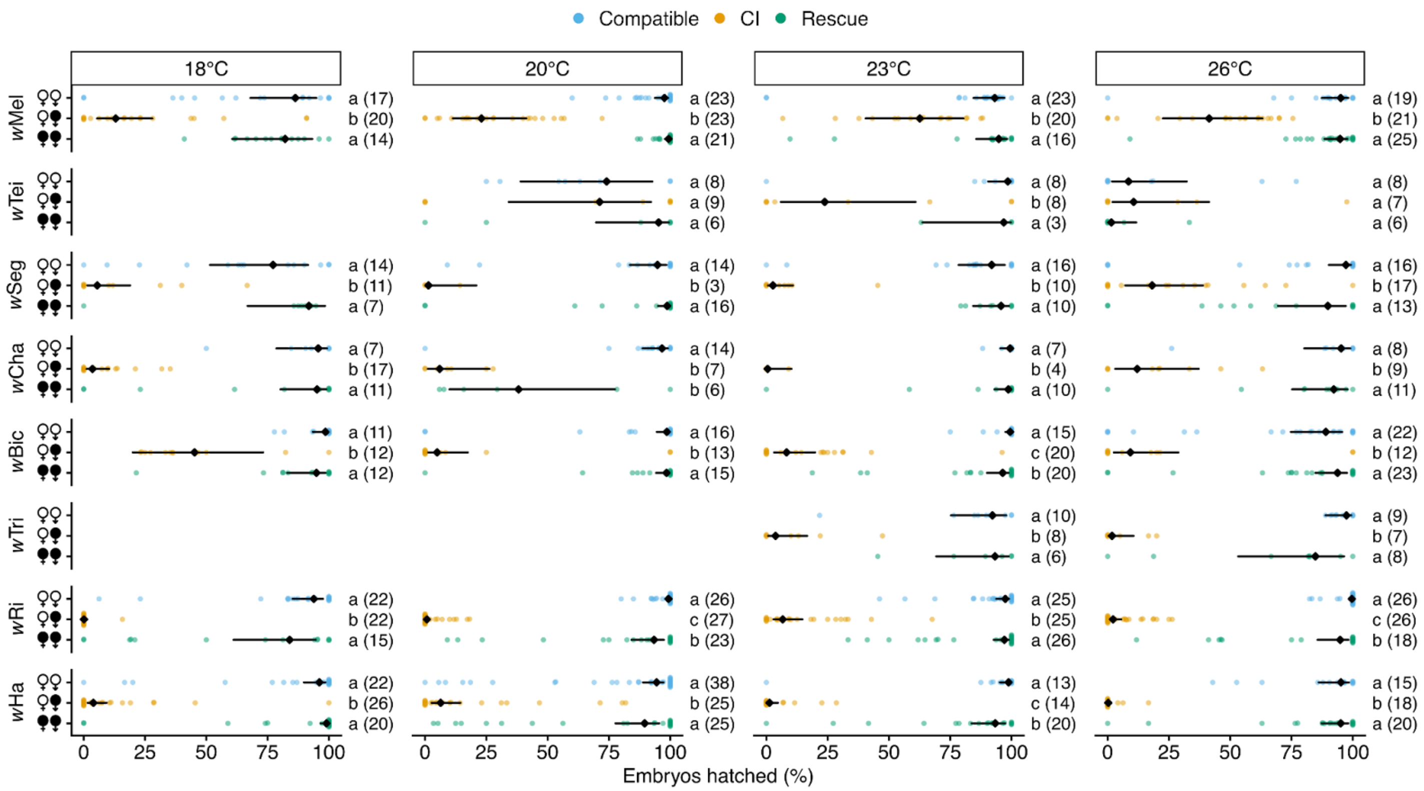


**Figure S4. CI crosses reduce embryonic hatching in seven of eight focal *Wolbachia* strains and rescue is occasionally incomplete.** Embryonic hatch rates from mating pairs. Column facets represent temperatures (18°C, 20°C, 23°C, 26°C). Row facets represent *Wolbachia* strain-host combinations. Three cross types are distinguished by female (♀) and male (♂) symbols: filled symbols indicate *Wolbachia*-bearing (symbiotic) individuals while unfilled symbols indicate *Wolbachia*-free (aposymbiotic) individuals. Individual data points (blue, compatible cross; yellow, CI cross; green, rescue cross) show the percentage of embryos that hatched from each cross. Black diamonds represent estimated means with error bars depicting 95% confidence intervals from a zero-inflated binomial generalized linear mixed model. Letters indicate significant differences in CI strength observed among temperatures within each strain (FDR-corrected, α = 0.05); shared letters denote non-significant differences. Sample sizes per group are in parentheses.

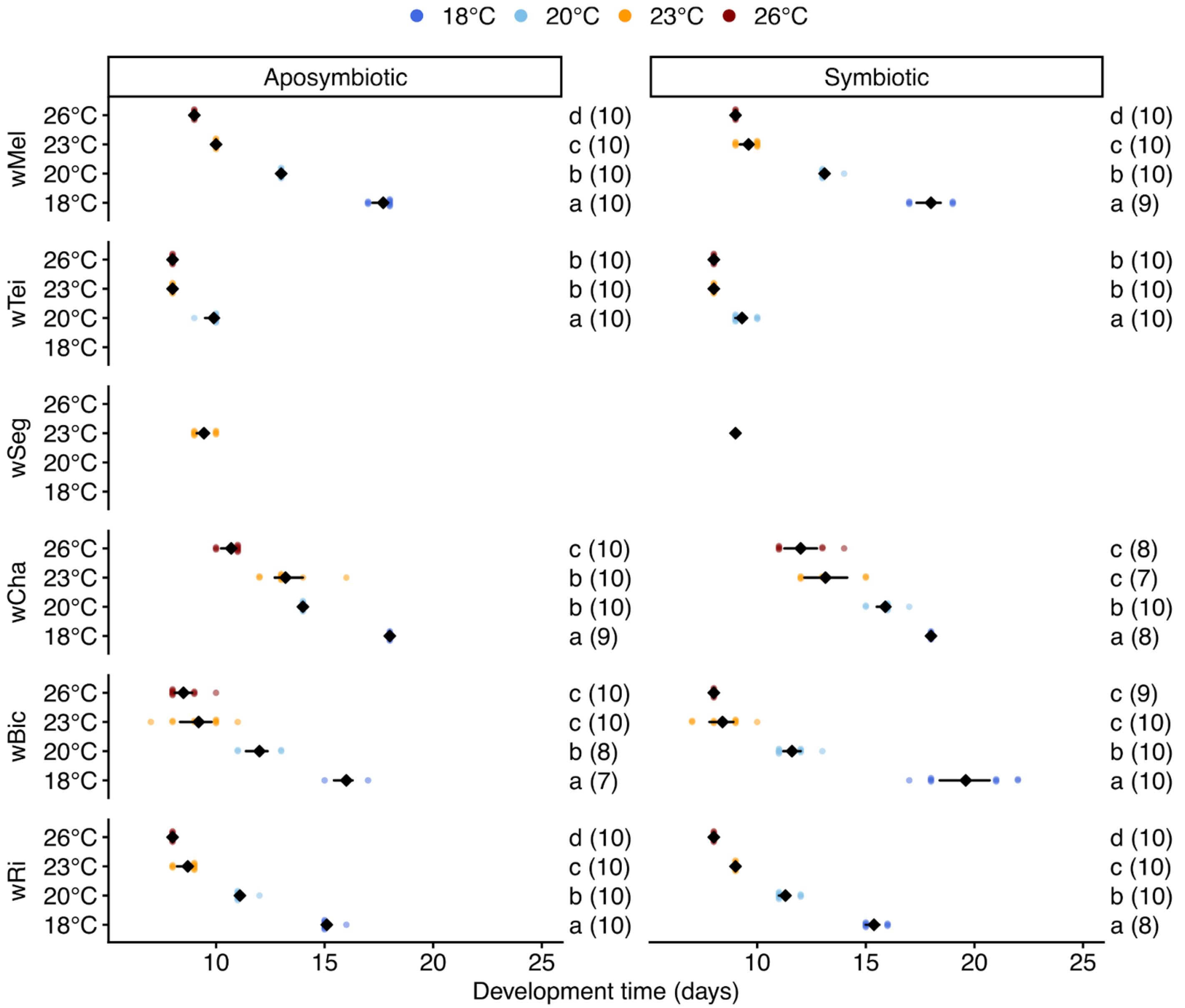


**Figure S5. Development time varies with temperature across all *Wolbachia*-host systems.** Egg-to-adult development time was measured across temperatures. Column facets represent cytotypes. Row facets represent *Wolbachia* strain-host combinations. Y-axis shows experimental temperatures (18°C, 20°C, 23°C, 26°C). Development time was measured as the number of days from egg laying to emergence of the first adult fly in each replicate vial. Individual data points show development time for each biological replicate, with blue dots representing aposymbiotic fly vials and green dots representing symbiotic fly vials. Black diamonds represent means with horizontal error bars depicting 95% bootstrap confidence intervals. Error bars are shown only for groups with within-group variance; groups where all observations were identical display diamonds without error bars. Letters indicate significant differences in CI strength observed among temperatures within each strain from pairwise permutation tests with 10,000 permutations (FDR-corrected, α = 0.05); shared letters denote non-significant differences. Sample sizes per group are in parentheses.

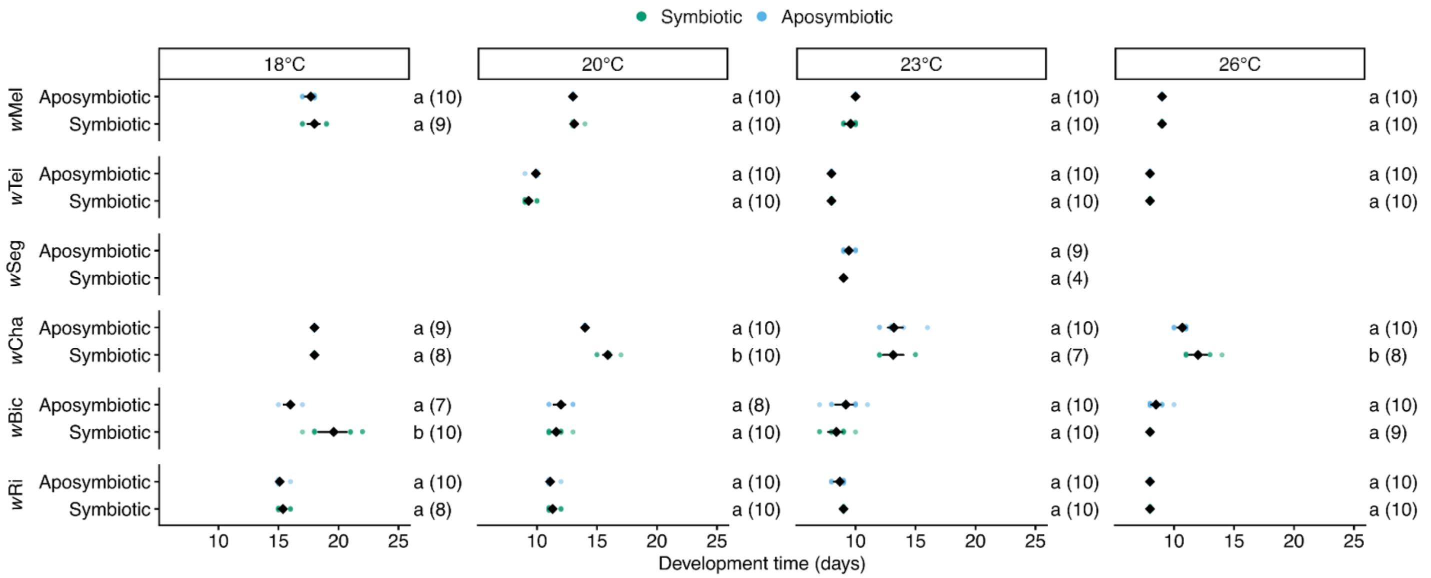


**Figure S6. *Wolbachia* accelerates development time in *w*Cha- and *w*Bic-bearing flies at specific temperatures.** Egg-to-adult development time was measured across cytotypes, host systems, and temperatures. Column facets represent experimental temperatures (18°C, 20°C, 23°C, 26°C). Row facets represent *Wolbachia* strain-host combinations. We measured development time as the number of days from egg laying to emergence of the first adult fly in each replicate vial. Individual data points show development time for each biological replicate, with blue dots representing aposymbiotic fly vials and green dots representing symbiotic fly vials. Black diamonds represent means with horizontal error bars depicting 95% bootstrap confidence intervals. Error bars are shown only for groups with within-group variance; groups where all observations were identical display diamonds without error bars. Letters indicate significant differences in CI strength observed among temperatures within each strain from pairwise permutation tests with 10,000 permutations (FDR-corrected, $\alpha = 0.05$); shared letters denote non-significant differences. Sample sizes per group are in parentheses.

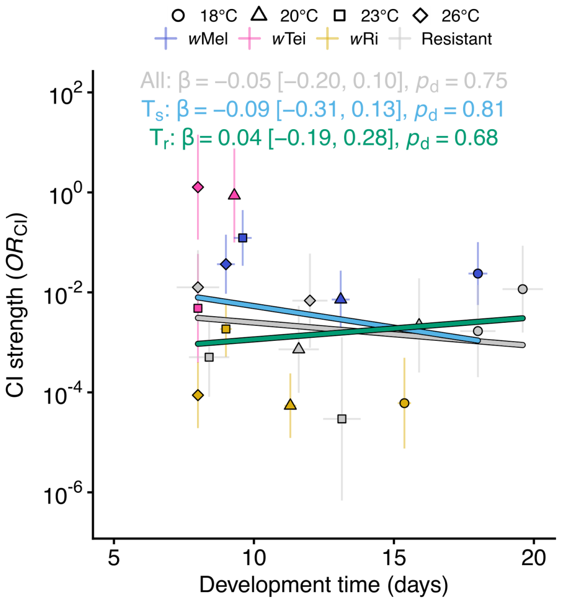


**Figure S7. CI strength does not correlate with development time.** Relationship between CI strength and development time across strain-temperature combinations. Each point represents estimated marginal means for a strain-temperature combination: CI strength ($OR_{CI}$) derived from a binomial mixed model of egg hatch, and development time from per-strain linear models of days to first adult emergence. Point shape indicates temperature (circle = 18°C, triangle = 20°C, square = 23°C, diamond = 26°C) and color indicates strain identity: strains with temperature-sensitive CI ($T_s$; *w*Mel = blue, *w*Tei = pink, *w*Ri = yellow) and strains with temperature-resistant CI ($T_r$; *w*Cha, *w*Bic = gray). Error bars indicate 95% confidence intervals for each estimated marginal mean estimate. Regression lines and corresponding statistics (β, 95% credible interval, $p_d$) are derived from Bayesian phylogenetic mixed models fit separately to all strains (gray), $T_s$ (light blue), and $T_r$ (green). Sample sizes: All (*n* = 19, 5 strains), $T_s$ (*n* = 11, 3 strains), $T_r$ (*n* = 8, 2 strains).

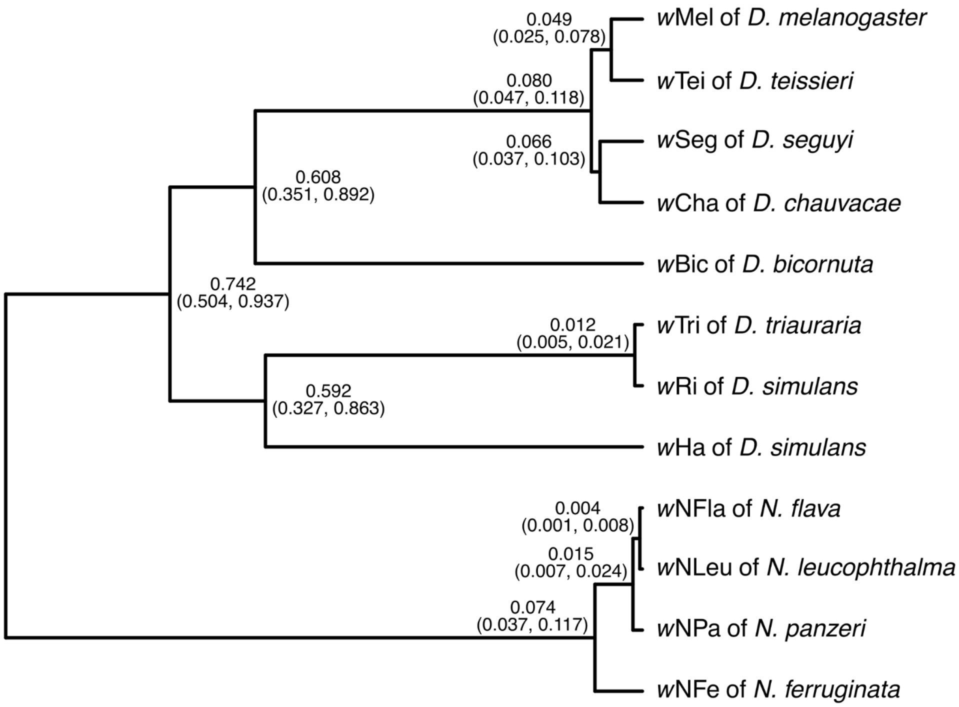


**Figure S8. Relative chronogram of the focal *Drosophila-associated Wolbachia*.** A relative chronogram based on 331 genes (285,540 bp). Four *Wolbachia* [(((*w*Nfla,*w*Nleu),wNpa),*w*Nfe)] from *Nomada* bee species were included as an outgroup to the eight *Drosophila*-associated *Wolbachia*. All *Wolbachia* belong to supergroup A and include closely related *w*Mel-like (*w*Mel, *w*Tei, *w*Seg, and *w*Cha) and *w*Ri-like (*w*Tri and *w*Ri) *Wolbachia*, as well as *w*Bic and *w*Ha. The eight *Drosophila*-associated variants diverged approximately 1.4 to 22 MYA according to the estimates from Shropshire et al. (2026). The root age was fixed to 1, and all nodes had posterior probabilities of 1. Nodes are labeled with the point estimates and 95% confidence intervals for relative age. See **Table S1** for the genome accessions.
